# Central T3 deprivation disturbs cortical cilia formation, oligodendrocyte lineage and neuronal cell-cell-communication in a MCT8/OATP1C1 deficient Allan-Herndon-Dudley Syndrome mouse model

**DOI:** 10.64898/2025.12.23.696195

**Authors:** Anna Molenaar, Ekta Pathak, Miriam Bernecker, Noémi Mallet, Ruth Gutierrez-Aguilar, Cristina Mencias, Dominik Lutter, Meri De Angelis, Gandhari Maity, Peter Kühnen, Benedikt Obermayer, Robert Opitz, Sonja C. Schriever, Timo D. Müller, Paul T. Pfluger

**Author notes:** Equal contribution (Anna Molenaar, Ekta Pathak). Corresponding author Prof. Dr. Paul T. Pfluger, Research Unit NeuroBiology of Diabetes Institute for Diabetes and Obesity Helmholtz Munich, Ingolstaedter Landstr.1 D-85764 Neuherberg.

## Abstract

**Background:** The Allan-Herndon-Dudley syndrome (AHDS) is a rare, X-linked human genetic disorder caused by mutations in the monocarboxylate transporter 8 (MCT8), essential for thyroid hormone (TH) transport across the blood-brain-barrier. The resulting central TH deprivation disrupts brain maturation and function, leading to intellectual disability and movement disorders. Cortical development, highly dependent on TH, is particularly affected and contributes significantly to AHDS pathologies.

**Methods:** To elucidate disrupted cortical processes, we conducted single nucleus RNA sequencing in a mouse model engineered to mimic the central TH deficiency characteristic of human AHDS. This murine AHDS model features the concomitant deletion of both MCT8 and OATP1C1, a T4 transporter largely absent in human brain capillaries that in mice plays a role in TH transport. The phenotype of dKO mice bears striking resemblance to the pathologies observed in human AHDS patients.

**Results:** Single nuclei were isolated from the cortex and attached striatum of 21-day old WT and MCT8/OATP1C1 dKO mice and sequenced using the 10x Genomics workflow. Cell proportion analyses on the resulting 48 clusters suggested elevated numbers of GABAergic striatal D1 and D2 neurons in the dKO mice. Diminished levels of mature oligodendrocytes coincided with a bifurcation within the oligodendrocyte lineage trajectory, leading to distinct subpopulations of WT and dKO oligodendrocytes. Differentially expressed gene (DEG) patterns align poorly with *Slc16a2* and *Slco1c1* mRNA levels in the respective clusters, but closely with prior published cortical bulk RNAseq data of mice with systemic hypothyroidism or MCT8/OATP1C1 deficiency. These parallels confirm the reliability of our data and provide new insights by pinpointing TH-responsive DEGs to specific cellular clusters. Moreover, inferred cell-cell communication using NeuronChat suggested a disbalance in GABAergic versus glutamatergic signaling. We further uncovered perturbed primary cilia formation in several GABAergic and glutamatergic clusters of the dKO cortex.

**Discussion:** Molecular signatures and perturbations uncovered by our snRNAseq study reveal new molecular characteristics of the AHDS. The imbalance in GABAergic versus glutamatergic cell-cell-communication, perturbed primary cilia formation, and bifurcation of the oligodendrocyte lineage align with pathologies observed in AHDS patients and highlight the role of TH signaling in maintaining neuronal network homeostasis in the cortex.

## Introduction

The Allan-Herndon-Dudley Syndrome (AHDS) is a rare but severe neurological disorder caused by genetic deficiency of the monocarboxylate transporter 8 (MCT8) [1], a highly specific transporter of triiodothyronine (T3), thyroxine (T4), as well as the inactive reverse T3 (rT3) [2]. The absence of MCT8-mediated transport of these thyroid hormones (TH) to the brain results in profound developmental, cognitive, and neuromotor deficits [3][4][5], as TH are involved in neurogenesis, gliogenesis, neuronal differentiation, and the maturation of synaptic connections [6][7][8][9]. Disruptions in these processes cause hypomyelination, delayed development of the cerebral cortex and cerebellum with aberrant cortical layering, as well as disruptions in parvalbumin (Pvalb) and neurofilament light chain expression in AHDS patients, already prenatally [10].

MCT8 is widely expressed in the developing brain and found in cells ranging from fetal cortical neurons, astrocytes, leptomeningeal cells, radial glia, choroid plexus epithelial cells, tanycytes to endothelial cells [11]. Cells of the blood brain barrier (BBB) or the blood cerebrospinal fluid barrier (BCSFB), like many other TH target cells in the brain, express also other TH transporters, contributing to dynamic and complex TH transport kinetics that remain incompletely understood. However, the severe symptoms caused by deficiency of MCT8 highlight that this TH transporter is of particular importance for CNS developmental processes.

In mice, studies on pathophysiological mechanisms underlying AHDS and molecular pathways affected by brain T3 deprivation had been complicated by the presence of an alternative transporter, OATP1C1, which may partially compensate for the lack of MCT8 to mitigate the severity of the phenotype observed in humans with AHDS [12]. OATP1C1 delivers T4 across the BBB of rodents but not humans, which lack expression in micro-vessels [13]. To better mimic the human condition, a double-knockout (dKO) mouse model deficient in both MCT8 and OATP1C1 has been developed, that exhibits symptoms and molecular pathologies akin to those observed in AHDS; these include brain TH levels, alterations in myelination and in the number and functionality of GABAergic Pvalb interneurons as well as motor dysfunctions presenting in gait abnormalities, albeit with less severity compared to the human syndrome [12][14].

Despite the insights gained from these models, a comprehensive characterization of the dKO’s effect on different cortical cell types remains elusive. This gap in knowledge has prompted us to initiate further investigation into the molecular impacts of TH deprivation, particularly during critical periods of postnatal cortical development. For this purpose, we applied single nucleus RNA sequencing (snRNAseq) to dissect the cellular and molecular consequences of MCT8 and OATP1C1 deletion in male wild-type (WT) and dKO mice after reaching an age of 21 days, a stage in which neurogenesis and myelination have been established. This approach allowed us to delineate the specific molecular alterations induced by TH deficiency, providing unprecedented insight into the hormone’s role in shaping the CNS landscape during early postnatal life.

## Results

Mice deficient for the TH transporters MCT8 and OATP1C1 (dKO) and the corresponding WT controls were sacrificed at day P21, when AHDS-specific pathologies such as neurogenesis and myelination have manifested (Fig. 1A). As described for adult mice [12][14], body weights of dKO mice at P21 (Fig. 1B) were significantly lighter than WT mice, even with comparable litter sizes (Fig. 1C) and the P21 dKO mice had increased plasma tT3, reduced plasma tT4 and higher tT3 to tT4 ratios, and profoundly reduced cortical tT3 and tT4 levels (Fig. 1 D-H). Brain weights, as well as the mass of individual brain areas, were not significantly different between genotypes (Fig. 1 I-J). Consistent with decreased TH levels in the cortex, we observed diminished mRNA levels of TH targets *Hr*, *Pvalb*, and *Aldh1a1* in dKO mice, and elevated levels of *Dio2*, whereas gene expression levels of *Gad1*, *Mbp,* and *Ald1l1* expression were not differentially regulated. Immunofluorescent staining for MBP revealed reduced protein amounts and a thinner corpus callosum (CC) in dKO mice (Fig. 1 L-N, Supplementary File 1).

**Figure 1.**
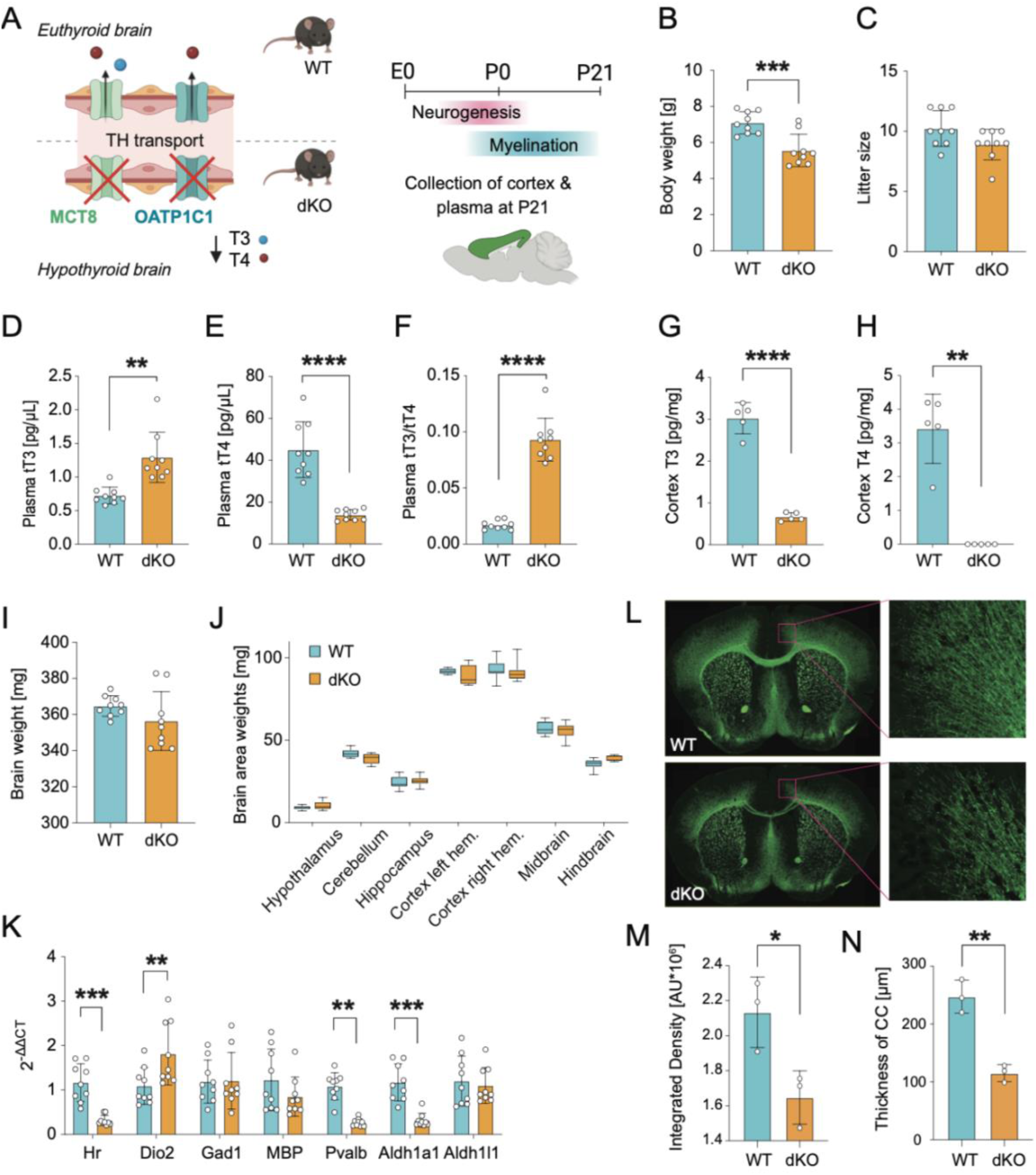
MCT8/OATP1C1 dKO mice mimic pathophysiologies of the Allan-Herndon-Dudley Syndrome. (A) The brains of dKO mice are hypothyroid due to the absence of MCT8 and OATP1C1 mediated thyroid hormone transport. Postnatal day 21 was chosen as endpoint to capture preceding changes in neurogenesis and myelination. At this age, dKO mice show decreased body weights (B), which are independent from (C) litter sizes. Circulating levels of total T3 (D), total T4 (E), and the T3/T4 ratio (F) in 21-day-old male dKO compared to WT mice, and total T3 (G) and T4 (H) in the brain are abnormal. Comparable weights for (I) whole brains and (J) specific brain regions at P21. (K) Cortical gene expression in WT and dKO mice at P21, showing differential expression of thyroid hormone-responsive and myelination-associated genes. (L) Immunohistochemical assessments for myelin basic protein (MBP) confirm an impaired myelination status in dKO mice, as seen by lower intensity (M), and thinner corpus callosum (N). Values represent means ± SD. (B-F, I-K) N = 9, (G-H) N = 5, (M-N) N = 3 (males per genotype). Statistical significance was determined using two-tailed unpaired Students’ t-tests with Welch’s correction (B-J), or multiple Students’ t-tests with the Holm-Sidak method (K). * p < 0.05, ** p < 0.01, *** p < 0.001, and **** p < 0.0001.

### Single nucleus RNA sequencing of WT and MCT8/OATP1C1 dKO mouse cortices reveal changes in cellular composition

To characterize cell-specific molecular events in the cortex that drive cognitive and locomotor impairments of dKO mice, we performed snRNAseq as outlined in Fig. 2A. The cortical hemispheres and attached cerebral nuclei of nine P21 male mice per genotype were isolated and then pooled into three nuclei suspensions for sorting. Fluorescence-assisted nucleus sorting (FANS) ensured a high-quality single nucleus suspension for GEM formation and sequencing using the 10x Genomics platform. After quality control steps (Suppl. Fig. 1A-C), dimensional reduction over all samples led to distinct cell types that were identified using MapMyCells [16], a tool provided by the Allen Brain Institute, as well as by highly expressed marker genes (Suppl. Fig. 1D). A total of 77897 nuclei were grouped into 48 clusters with 19 glutamatergic neuronal subtypes, 20 GABAergic neuronal subtypes, and 9 non-neuronal clusters subdivided into glial and vascular cell types (Fig. 2B). Using MapMyCells’ class annotation, glutamatergic and GABAergic neurons could be assigned to overarching groups of neurons belonging to the cerebral cortex (CTX), cerebral nuclei (CNU), or the lateral septum (LSX) and olfactory bulb (OB) (Fig. 2B).

**Figure 2:**
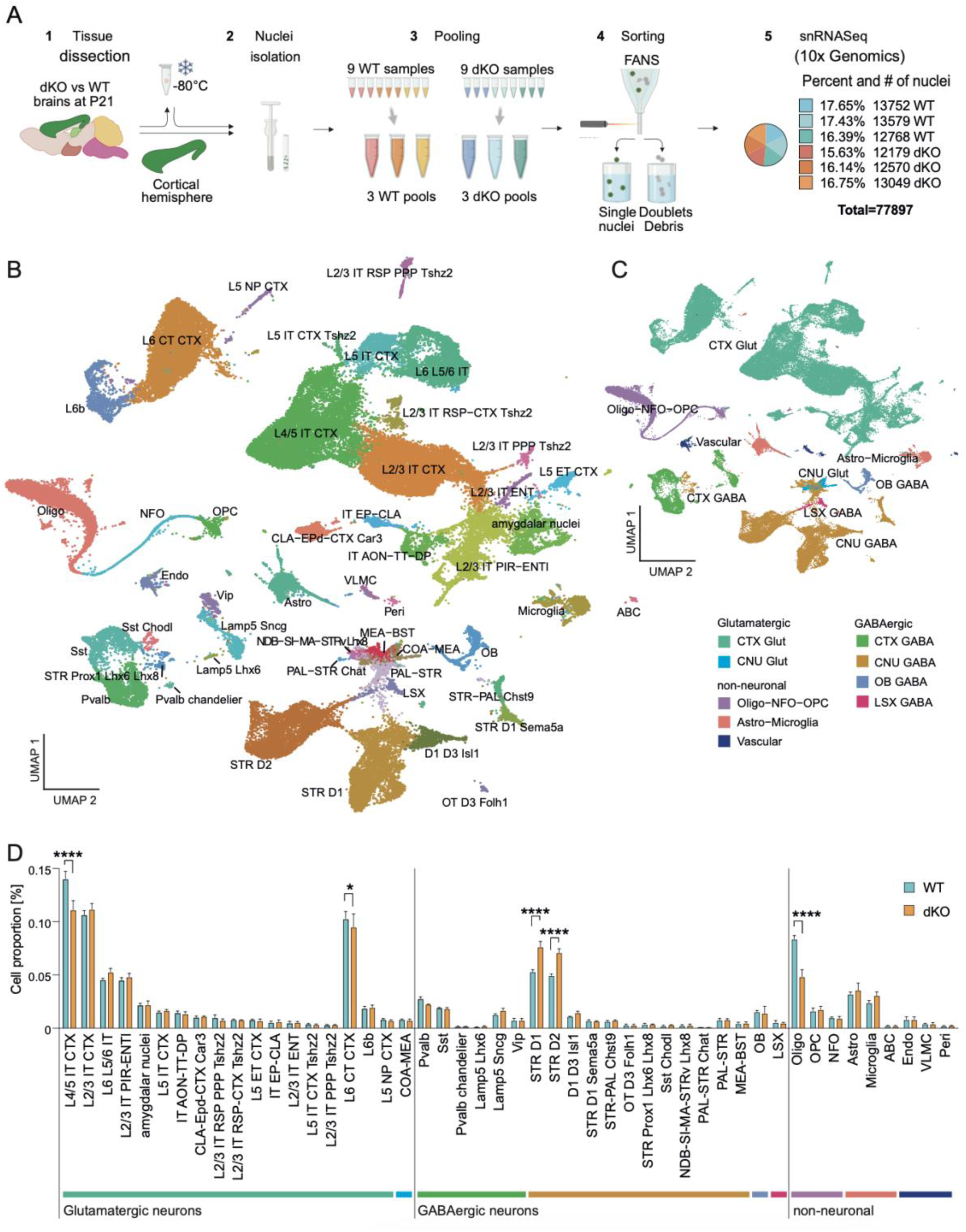
Single nucleus RNA sequencing of MCT8/OATP1C1 dKO mouse cortices. (A) Schematic of the snRNAseq pipeline for 3x 3 WT and dKO mouse cortices (male, age P21) that resulted in 77897 nuclei, with homogenous distribution between the WT and dKO samples. (B) UMAP depicting the resulting 48 clusters and (C) the overarching cell group they belong to. The nuclei grouped into glutamatergic and GABAergic neurons, and non-neuronal cell types, further split into cortical areas (CTX) and cerebral nuclei (CNU), as well as olfactory bulb (OB) and lateral septum (LSX), and glial and vascular cell types. (D) Relative cell proportions of the 19 glutamatergic, 20 GABAergic, and 9 non-neuronal cell types within each cluster between WT and dKO. Values represent means ± SD. N = 3 per genotype. Statistical significance was determined using multiple t-tests with Holm-Sidak correction. * p < 0.05, **** p < 0.0001.

Cell proportion analyses of our 48 clusters revealed elevated numbers of GABAergic striatal (STR) D1 and STR D2 neurons and diminished numbers of glutamatergic L4/5 IT CTX neurons in dKO mice (Fig. 2D), which could lead to a disbalance of excitatory and inhibitory signaling between the cortex and striatum. Consistent with the decrease of cortical myelination in the dKO mice, we found diminished levels of oligodendrocytes.

Differences in cell propensities were partially confirmed by immunofluorescence staining in P21 cortical slices. Olig2 staining for the identification of oligodendrocytes showed similar glial cell cumbers for both genotypes when comparing the entire cortex area, but reduced numbers in dKO mice when focusing the analysis to the corpus callosum (Fig. 2A). Using Rorb as a marker for L4/5 IT CTX cells, we could not find decreased cell numbers in dKO mice (Suppl. Fig. 2A) [17]. As Rorb antibodies could also stain astrocytes [18], we used RNAscope™ but again failed to identify a decrease in Rorb expression in neurons (Suppl. Fig. 2B). RNAscope™ for Pdyn and Penk as markers for STR D1 and D2 neurons confirmed an increase of these neurons in the striatum of dKO mouse brains at P21 (Fig. 2B). RNAscope further revealed a clear reduction in the number of cortical parvalbumin expressing Pvalb cells in dKO mice (Suppl. Fig. 2A), which are reported to be highly affected in dKO mice and AHDS patients [10][14][15][19].

### Differentially expressed genes in dKO vs WT mouse brains

Differential gene expression analyses on the overarching cell groups showed profound changes in CTX Glut, CTX GABA, and CNU GABA neurons, followed by the oligodendrocyte lineage (Suppl. Fig. 3A). The numbers of downregulated genes were slightly higher than upregulated genes in most cell groups (Suppl. Fig. 3B). In-depth analyses of the individual 48 clusters confirmed the trend toward downregulated genes. Highest numbers of DEGs were found for several glutamatergic neuronal clusters such as L4/5 IT CTX, L2/3 IT CTX, or L6 CT CTX, GABAergic clusters such as Pvalb, and the STR D1 and D2 neurons (Fig. 3A, Supplementary File 2). Of note, gene expression levels of *Slc16a2* (MCT8) and *Slco1c1* (OATP1C1) in our 48 WT clusters were poor predictors for the overall number of differentially expressed genes (DEGs) per cluster, indicating either developmental perturbations at earlier stages of development, or non-cell-autonomous effects due to the deprivation of central TH. To gain an understanding on processes affected in the brains of P21 dKO mice, we analyzed GO Biological Process terms that were enriched in the DEGs of the overarching groups using EnrichR (Fig. 3B, Supplementary File 3). As expected for brains with perturbed TH-availability, various processes of brain development as well as brain function such as synaptic signaling or ion transport were impaired in most neuron types of the cerebral cortex and cerebral nuclei as well as glial groups. In CNU Glut, LSX GABA, and OB GABA cells, we found an enrichment for biological processes related to the regulation of cell migration and mitochondrial membrane organization.

**Figure 3.**
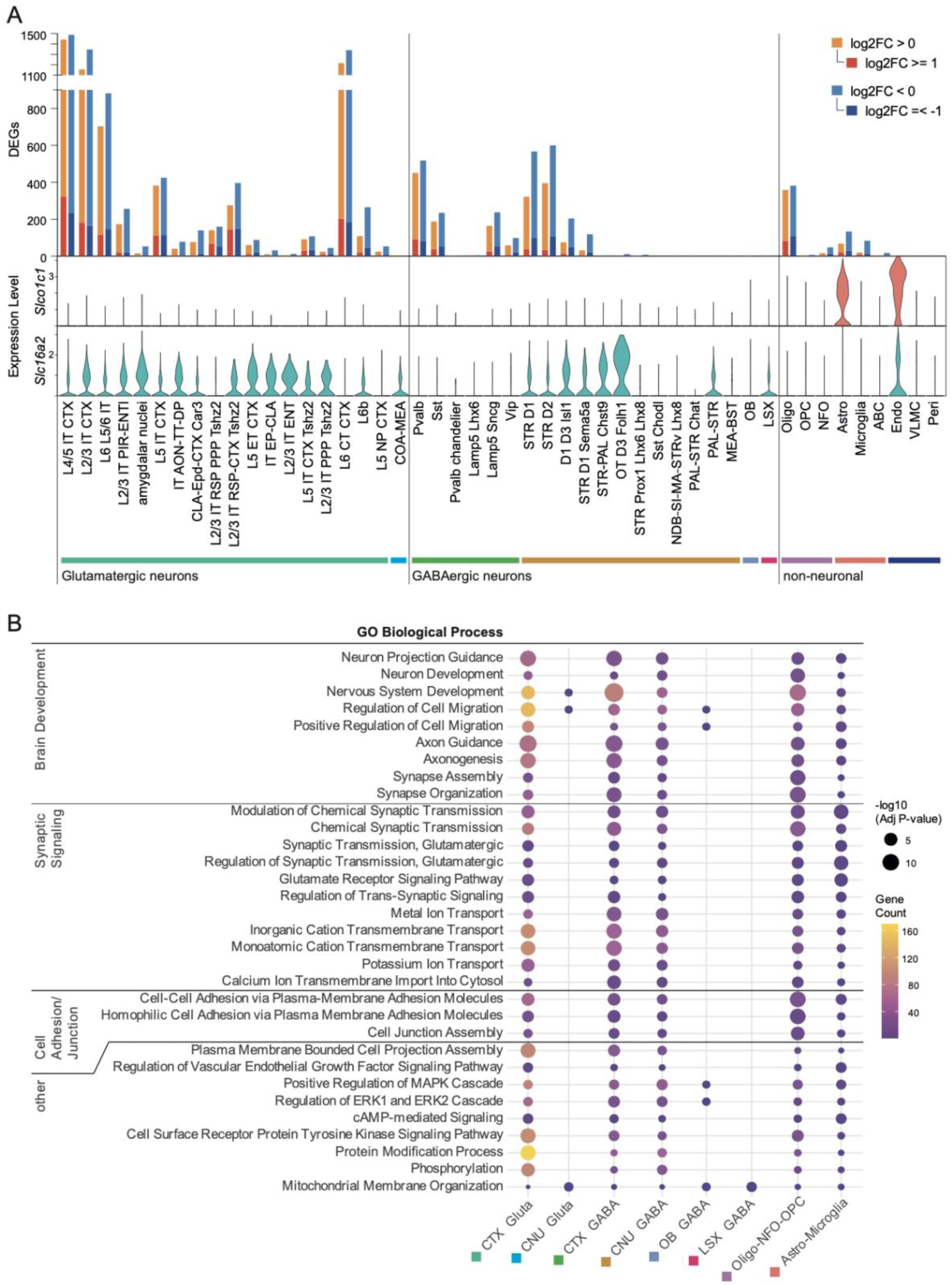
Gene expression changes in WT and dKO mouse cortices. (A) Differentially expressed genes (DEGs) per cluster with padj < 0.05, those upregulated in dKO in red/orange and downregulated in blue/light blue. Red and blue are genes that additionally have a log2FC greater than 1 or less than −1. Below are the expression levels of *Slco1c1* and *Slc16a2* in the WT portion of the clusters. (B) Enrichment of biological processes in the DEGs of the overarching groups using EnrichR [2].

### Cilia dysfunction in the cortex of dKO mice

To gain deeper insights into the biological patterns underlying the DEGs identified between WT and dKO clusters, we conducted gene set enrichment analysis (GSEA) (Supplementary File 4), plotting biological processes enriched in at least two clusters (Fig. 4A). With this method, many terms demonstrated enrichment that were associated with cilia structure and function, including the axoneme and microtubules (indicated in green). Neuronal cilia, which are primary cilia, are crucial signaling hubs during neurogenesis, neuronal migration, axon pathfinding, and cortical circuit formation [20]. Genes extracted from the respective GSEA terms and additional, published ciliopathy gene sets [21][22][23] were significantly deregulated in the main clusters showing the cilia gene set enrichments (Fig. 4B), most notably *Nckap5*, associated with microtubule bundle formation. Its upregulation might indicate an increase in cilia assembly in the dKO cortex. Staining for cilia marker Adcy3 (AC3) to confirm perturbations in cilia structure *in vivo* showed an increased number of cilia in dKO mice. In the somatosensory cortex the length of the cilia was significantly increased (Fig. 4 C,D).

**Figure 4:**
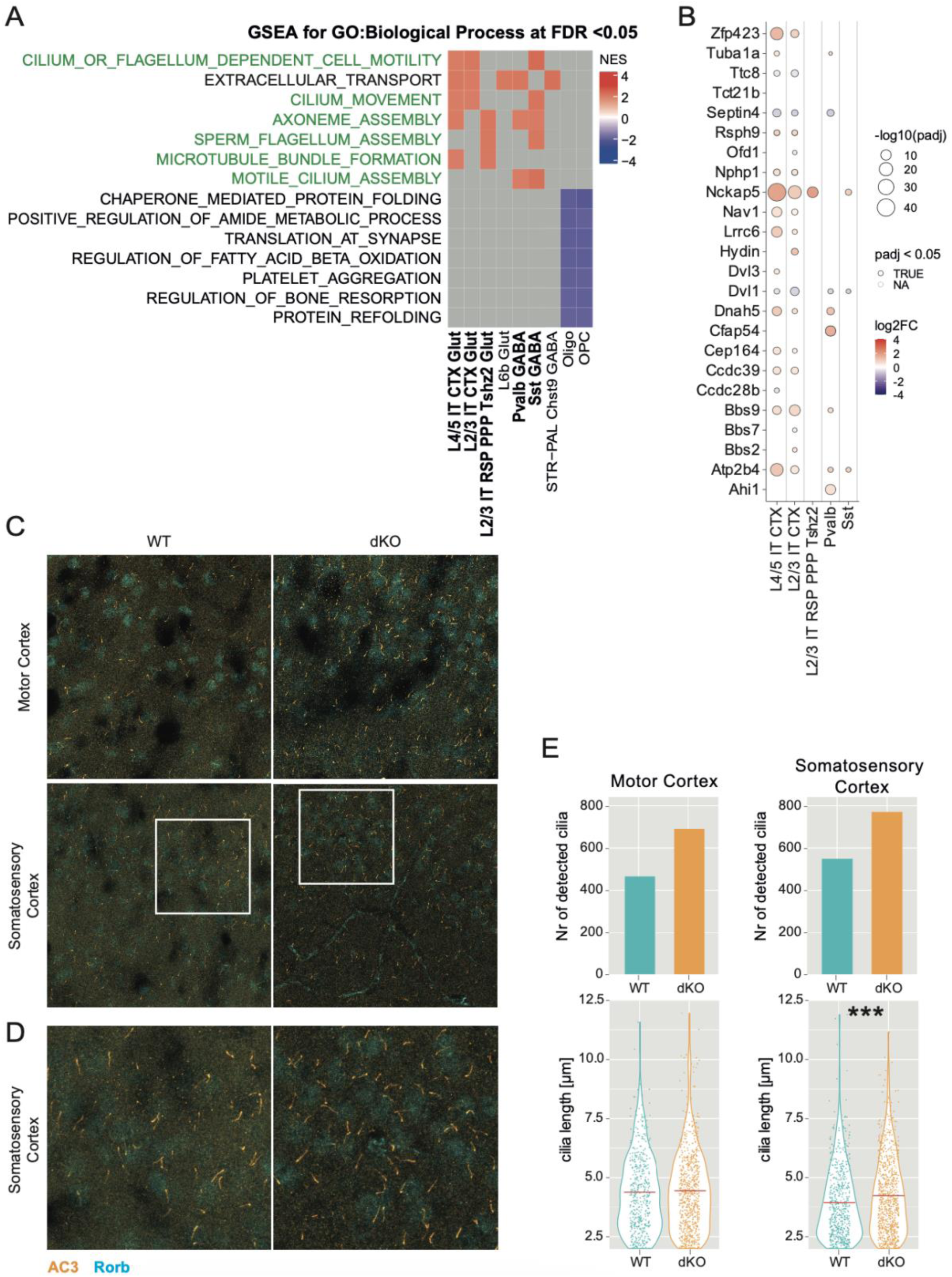
Gene Set Enrichment Analyses indicate cilia dysfunction. (A) GSEA for Biological Processes enriched or depleted in dKO cortices in at least two clusters. Terms related to cilia in GABAergic neurons and Glutamatergic are marked in green. NES: enrichment score normalized to mean enrichment of random samples of the same size (B) Dot plot for selected clusters from (A, bold) reveals differential expression of genes linked with impaired ciliogenesis and ciliopathologies in dKO mice. (C) Stainings in WT and dKO mouse motor cortex and somatosensory cortex using AC3 as cilia marker (orange), co-stained with Rorb (blue). (D) Zoom of area marked with white box in (C). (E) Cilia parameters derived from (C), using the entire field of view of the z-stack confocal images. Cilia analysis was done using CiliaQ [3]. N = 6 per genotype and area. Statistical significance was determined using two-tailed unpaired Students’ t-tests with Welch’s correction. *** p < 0.001. Mean marked with red line.

### Thyroid hormone related gene expression signature in the dKO cortex

Next, to contextualize our findings, we compared our snRNAseq-derived gene expression profiles with publicly available bulk RNA-seq data from systemic and central T3 deprivation models, assessing the relevance of known thyroid hormone-responsive markers in the dKO cortex. Specifically, we combined the cortical (CTX) clusters and the non-neuronal (NN) clusters as pseudobulk cortex, and the cerebral nuclei cells (CNU) as pseudobulk striatum, and compared those with published bulk seq data from a P21 mouse cortex and striatum of another MCT8/OATP1C1 dKO (MO) mouse model as well as a systemically hypothyroid (SH) mouse model [24] (Fig. 5A). Overall, we found striking directional correlations between the DEGs common between our dKO pseudobulk, SH and MO (Suppl. Fig. 4A). A Venn diagram of DEGs reveals both a substantial overlap between all three groups for cortex and striatum but also DEGs unique to the three conditions, likely stemming from the different sequencing techniques as well as slight differences in analyzed cell types (Fig. 5B, Supplementary File 5). Additionally, there are distinct DEGs found in our dKO pseudobulks and MO but not in SH, which could help describe effects specific to the KO of MCT8 and OATP1C1 as opposed to systemic hypothyroidism.

**Figure 5:**
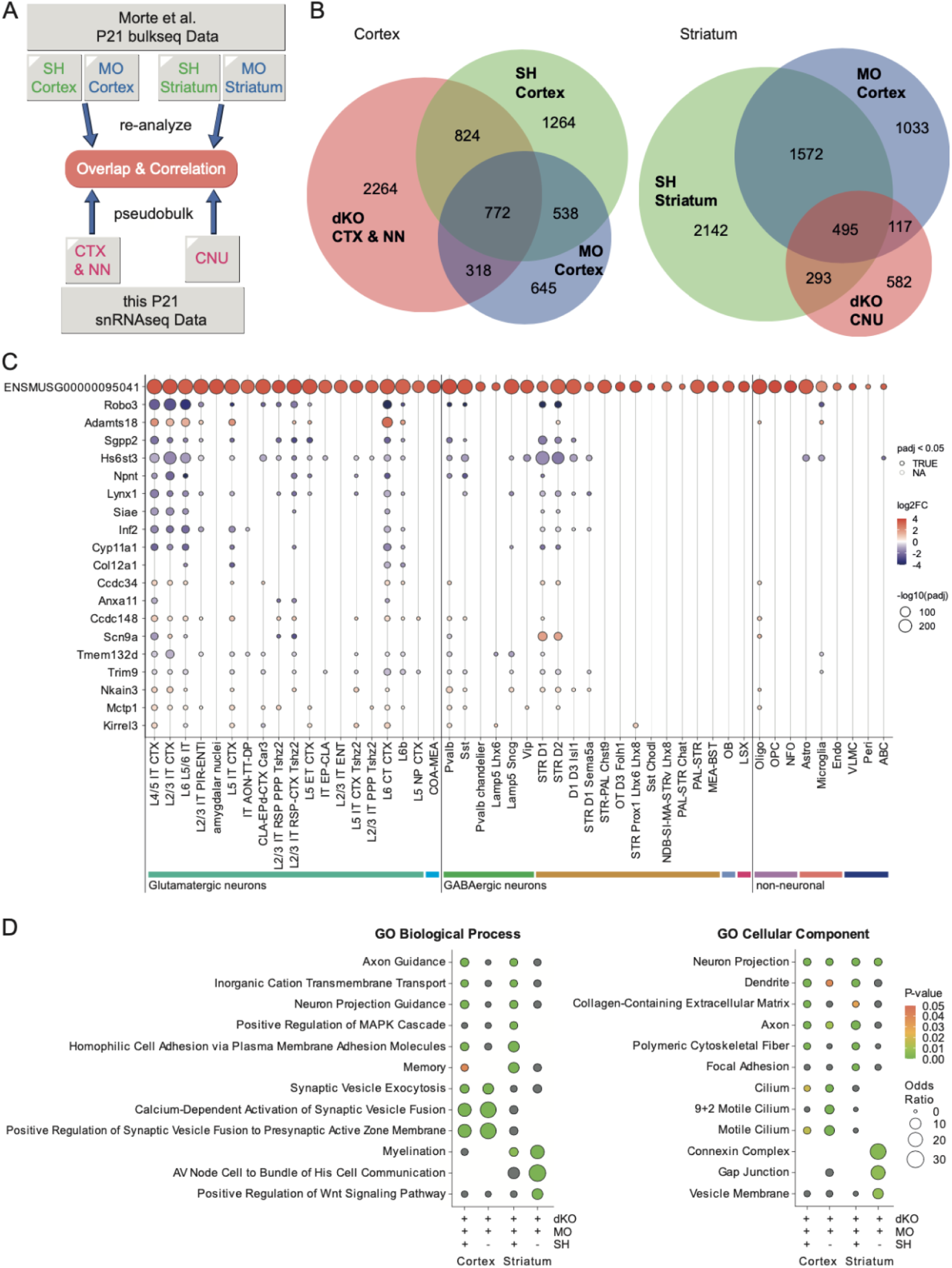
Correlations between our dKO cortical and striatal transcriptional signatures and public domain data of T3 deprivation models highlight shared but also distinct and region-specific pathway enrichments. (A) Bulkseq data from Morte et al. [4] of systemically hypothyroid (SH) mice and MCT8/OATP1C1 dKO (MO) mice at age P21 were re-analyzed and compared to the pseudobulk of our dataset. Cortical bulkseq data was compared to the combined cortical clusters (CTX) plus non-neuronal (NN) clusters, striatal bulkseq data was compared to the combined striatal clusters (CNU). (B) Venn diagram of the DEGs with padj < 0.05 from the MO and SH bulkseq data and the pseudobulk versions of our data for cortex and striatum. (C) Dotplot showing log2FC and padj per cluster in our single nucleus dataset of genes from the cortex comparison. Shown are genes either commonly differentially expressed in SH, MO and the pseudobulk of our dKO dataset, or commonly differentially expressed in MO and our dKO pseudubulk, but not in SH. (D) Enrichment of biological processes and cellular component in the DEGs overlapping in the cortex or striatum comparison of SH and MO and dKO, or of MO and dKO but not SH using Enrichr [5][2]. Grey means the term was found but is not significantly enriched, empty means the term has not been found. Input genes can be found in Supplementary Table 5. Venn diagrams generated using DeepVenn [6].

The cluster-specific plotting of dKO CTX & NN pseudobulk DEGs overlapping with both SH and MO cortex DEGs and sorted by their fold change mostly placed those within large glutamatergic clusters such as L4/5 IT CTX, L2/3 IT CTX, and L6 L5/6 IT, and GABAergic clusters such as Pvalb, STR D1 and STR D2 neurons (Fig. 5C). Among the downregulated genes were *Sgpp2* which is involved in migration, adhesion, survival, and proliferation [25], and *Robo3*, which is important for axon guidance [26] and was observed earlier to be T3-regulated in glutamatergic motor cortex neurons by Hochbaum et al. (2024) [27]. The metalloproteinase *Adamts18*, which modulates extracellular matrix and has been associated with grey matter integrity and cell-cell-adhesion [28], was particularly increased in glutamatergic clusters. One gene consistently upregulated in all dKO clusters was ENSMUSG00000095041. This gene is described as a putative pseudogene *(Pisd-ps3)* to *Pisd*, which encodes a phosphatidylserine decarboxylase. The upregulation in the P21 dKO brains was confirmed by qPCR analyses (Suppl. Fig. 4B). *Pisd-ps3* was also upregulated in the heart, while levels remained comparable for the liver and kidney. Among the genes deregulated in the dKO pseudobulk and MO but not SH are E3 ubiquitin ligase *Trim9* and synapse adhesion protein *Kirrel3*, both associated with synapse formation in the brain [29][30]. Commonly deregulated genes between dKO, SH, and MO were enriched in biological processes related to axon guidance and neuron projection guidance, whereas genes only common in dKO and MO but not SH where more significantly enriched in terms related to synaptic vesicle fusion and exocytosis for the cortex comparison, or in myelination and Wnt signaling for the striatum comparison. Cellular component terms were also differentially enriched, with neuron projection genes being enriched for the overlapping genes of all comparisons, cilium component genes enriched for the overlap of cortical dKO and MO DEGs without SH DEGs, and connexin complex and vesicle membrane genes being enriched for the DEGs of dKO and MO and not SH in the striatum (Fig.5 D).

Regarding the general state of TH signaling, known effectors of thyroid hormone action remained largely unaffected (Suppl. Fig. 4C-D). Thyroid hormone receptor *Thra* expression was upregulated and well-known TH target genes *Hr* and *Klf9* expression levels were downregulated in many dKO clusters, but the expression levels of co-regulators, deiodinases 2 and 3, or other TH transporters were mostly unchanged and/or barely detectable in WT and KO mice (Suppl. Fig. 4D)

### NeuronChat analyses delineate perturbations of cellular communication in the P21 dKO cortex and striatum

Given the observed alterations in cell type proportions, reduced myelination, and transcriptomic signatures pointing to disrupted axonal guidance and synaptic organization as well as transmission, we next explored whether cellular communication between glutamatergic and GABAergic clusters is perturbed by applying the NeuronChat package to infer cell-cell communication based on known ligand-receptor interactions [31]. Overall, we found slightly reduced interaction strengths but an increased number of inferred interactions between the overarching cell groups in dKO compared to WT mice (Suppl. Fig. 5A). Comparing the signal strengths of typical interaction pairs of inhibitory and excitatory signaling between WT and dKO unraveled changes in outgoing and incoming signaling between neuron cell groups; GABAergic outgoing signals were increased for CNU and LSX neurons (Fig. 6A) and incoming signal strengths were broadly increased for the GABAergic interaction pairs between all cell groups (Fig. 6B). Likewise, the number of GABAergic interaction links were profoundly elevated in dKO compared to WT mice (Suppl. Fig. 5B). Information flow for the interaction pairs Nrxn3-Nlgn1 and Nrx1-Nlgn1 was elevated in WT compared to dKO mice, while GABAergic interactions as well as Nrxn2 interactions with Nlgn1 had lower information flow in WT compared to dKO mice (Suppl. Fig. 5B). Altered neurexins (Nrx) - neuroligin (Nlgn) interactions have been implicated in perturbations of synapse formation and plasticity [32]. Outgoing glutamatergic signals were mostly reduced in dKO CTX Glut neurons, and incoming glutamatergic signals varied across cell groups in the dKO, with some showing slight increases and others showing decreases depending on the interaction pair.

**Figure 6:**
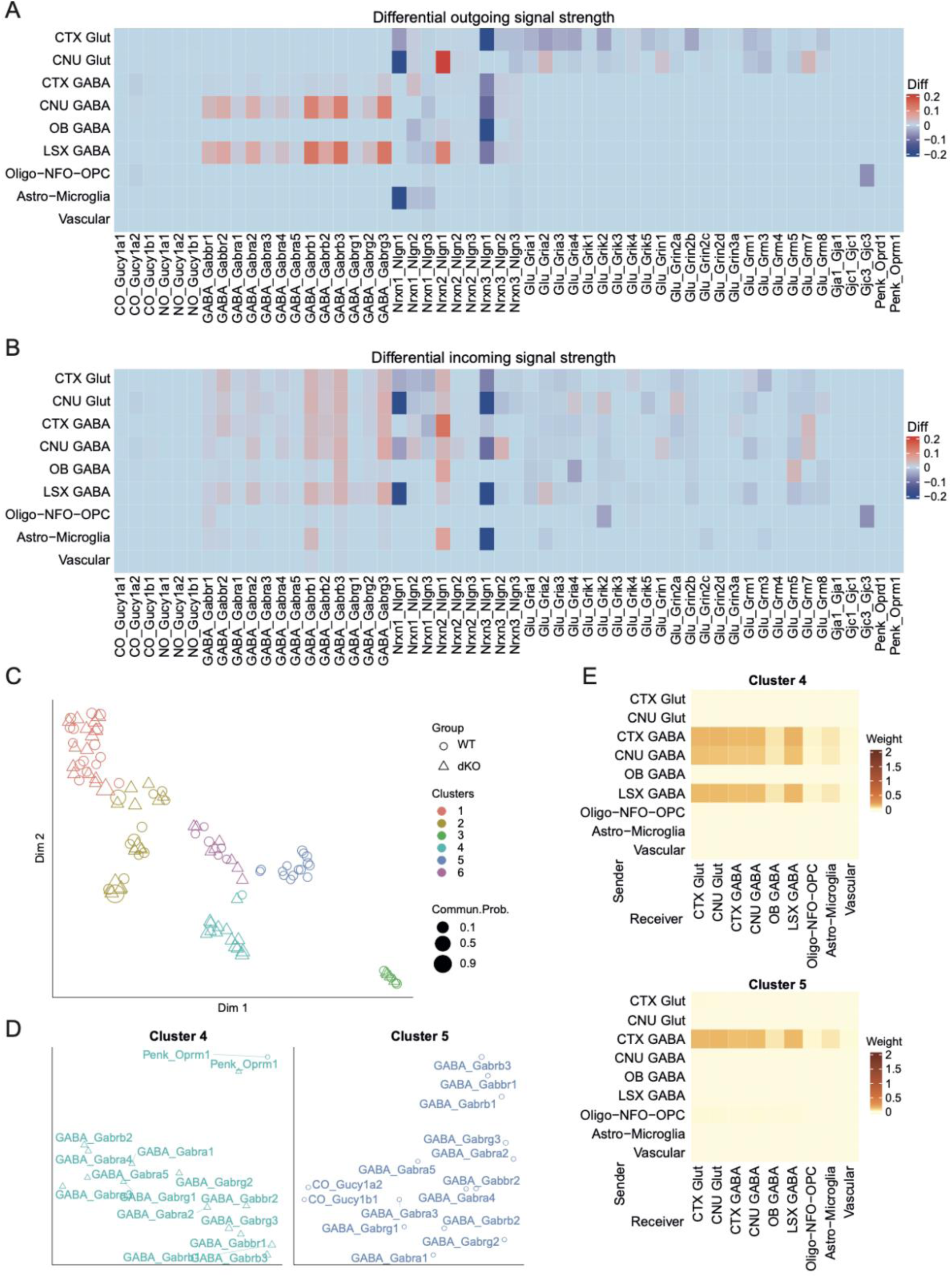
NeuronChat-based cell-cell-communication profiling reveals distinct GABAergic signaling patterns across cortical and subcortical regions in WT and dKO mice. Heatmaps of (A) differential outgoing and (B) differential incoming signal strength of specific interaction pairs (x-axis) for WT vs dKO mice between the overarching cell groups (y-axis) using NeuronChat [7]. (C) A two-dimensional manifold projection of the interaction pairs revealed five distinct clusters based on sender-receiver similarities, (D) Cluster 4 and cluster 5 show communication networks unique to dKO mice and WT mice respectively (see supplemental figure 5 for the other clusters). (E) Heatmaps depicting the weight/strength of communication signals between the overarching cell groups as senders and receivers for specific interaction-pair clusters 4 and 5. The respective weights of signal in the heatmaps represent the sum of communication strength values over all interaction pairs of that cluster. Statistical significance was assessed with NeuronChats integrated statistical tests, with p-values corrected for multiple tests using the Benjamini-Hochberg procedure.

After projecting the interaction pairs from WT and dKO onto a shared two-dimensional manifold and applying k-means(k=6) clustering to its first two components, we revealed that most interaction-pairs had similar senders and receivers per genotype, with similar communication probabilities (Fig. 6C; Suppl. Fig. 5C,D). However, multiple GABAergic interaction-pairs (Fig. 6D) clearly separated genotype-specific into distinct cluster 4 for dKO and cluster 5 for WT mice. Strongest communication strengths for dKO cluster 4 were thereby observed from the sending GABAergic neuron groups CTX, CNU, and LSX to most other GABAergic as well as glutamatergic neuron groups and to lesser extent glia, whereas in WT, only the CTX GABA group interacted with several receiving, predominantly neuronal groups (Fig. 6E). Cell-type specific NeuronChat analyses were fully consistent with the group-wise comparison and perturbed outgoing vs incoming signal strengths of GABAergic and glutamatergic cell types (Suppl. Fig. 6A,B). When comparing WT and dKO mice for their distinct GABAergic signal strengths originating from the individual cell clusters, we found additional increases in signal strength for the CNU-specific STR D2 and the LSX cell cluster in dKO mice (Suppl. Fig. 7A-C). Overall, our collective data point toward an increase in outgoing GABAergic interactions for CNU and LSX neuronal populations that may exert a wide-spread shift in the excitation/ inhibition (E/I) balance in the dKO mice.

### Altered oligodendrocyte maturation pathways and their potential impact on myelination in dKO mice

Neuronal imbalance can arise from various causes, including impairments in myelination and oligodendrocyte numbers, as observed in our dKO mice and in AHDS patients. To further delineate these alterations, we performed a sub-cluster analysis on the oligodendrocyte lineage of OPCs, NFOs and Oligo, identifying 8 clusters (Fig. 7A) with distinct marker genes (Fig. 7B). The trajectory analysis using Monocle3 (Fig. 7C) revealed a clear branching along the occurrence of mature oligodendrocytes (cluster 5), with a bifurcation of WT and dKO oligodendrocytes along distinct trajectories. Accordingly, clusters 1 and 3 were only populated by WT, and cluster 2 only by dKO oligodendrocytes (Fig. 7D). Overall, dKO mice had lower numbers of mature oligodendrocytes, but higher numbers of NFO in cluster 6 and OPCs in cluster 4, suggesting that oligodendrocytes may be impaired to progress from immature stages. While we highlighted impairments in myelination *in vivo* with MBP staining (Fig. 1L) and enrichment of myelination related genes in the overlap of striatal dKO and MO but not SH DEGs (Fig. 5D), prominent marker genes of myelination, oligodendrogenesis, or cellular communication demonstrated comparable expression levels in mature oligodendrocytes of dKO cluster 2 and WT clusters 1 and 3 (Fig. 7E). However, genes were enriched differentially for dKO and WT clusters along the bifurcation (Fig. 7 F,G), notably, cell adhesion molecules and genes related to the cytoskeleton and ECM (*Cdh13, Thsd4, Fnbp1, Kif13b, Magi2, Ank2, Cntn2, Cdh20, Lama2*), ion channels (*Kcna1*, *Kctd3*), axon guidance (*Dcc*), and *Bace2* and *Sirt2*, both associated with neurodegenerative diseases and aging [33][34]. Last, dKO oligodendrocytes had higher expression levels of lipopolysaccharide-induced TNF-alpha factor (*Litaf*), a signal transducer gene not yet linked with the AHDS but associated with the demyelinating Charcot-Marie-Tooth disease, which includes ataxia and distal muscle weakness [35], all three prominent symptoms of AHDS. By using RNAscope™ for the two genes with the clearest distinction in WT vs dKO enrichment, we confirmed the pattern at the cellular level, with Litaf being markedly upregulated in oligodendrocytes of dKO mice, whereas Lama2, a gene associated with congenital muscular dystrophy, weakened ECM and integrin signaling, and impaired structural and trophic support [36], was markedly higher in WT and down in dKO oligodendrocytes (Suppl. Fig. 8D-E). Collectively, these data are consistent with disturbed myelination and aberrant oligodendrocyte maturation and the transition toward a stress-associated, demyelination-prone state.

**Figure 7:**
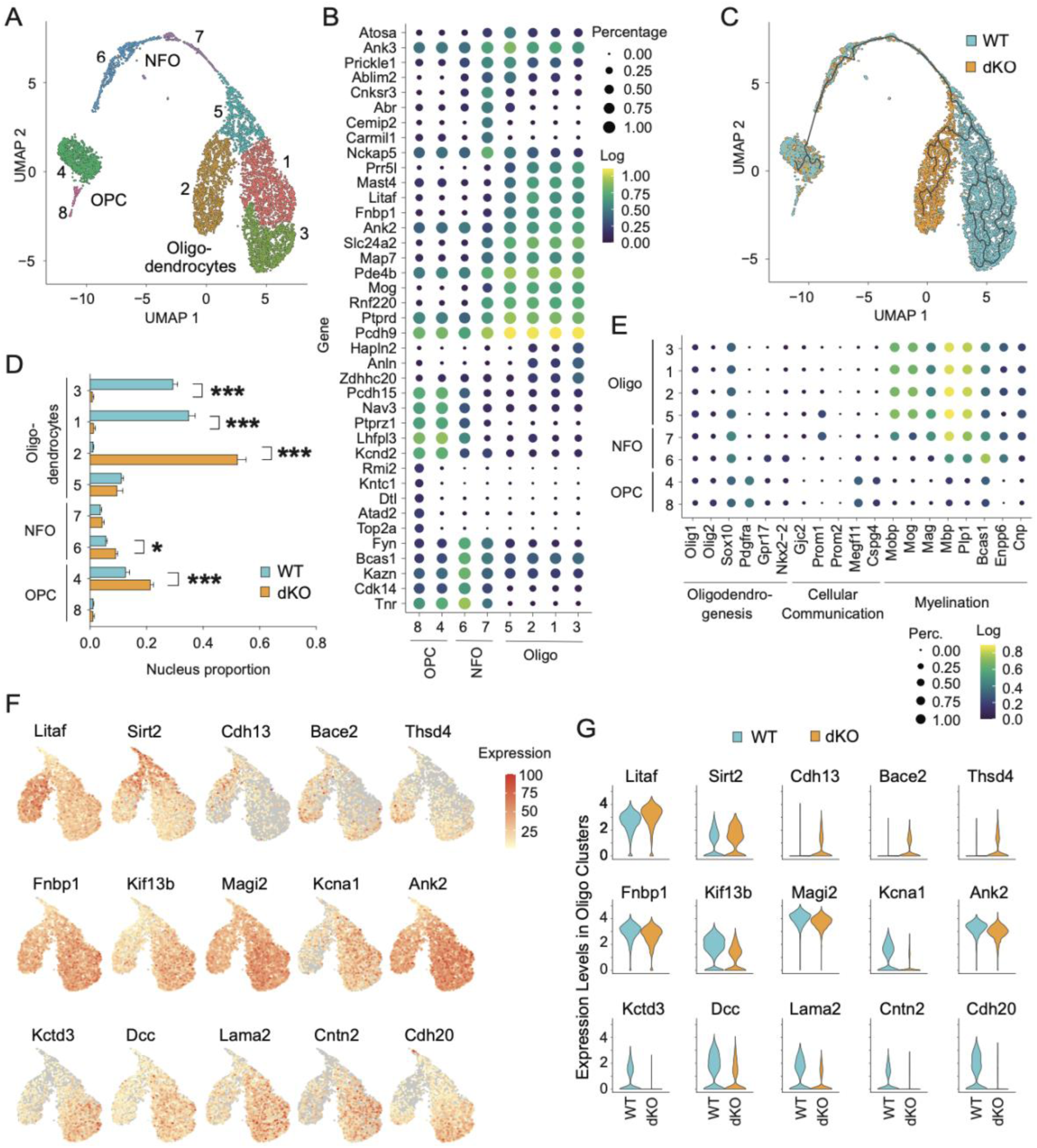
Perturbed oligodendrocyte lineage dynamics in WT and dKO cortices. Sub-clustering within the oligodendrocyte cell lineage is visualized by the (A) UMAP of distinct oligodendrocyte precursor cell (OPC) clusters, newly formed oligodendrocyte (NFO) clusters and mature oligodendrocyte clusters. (B) Dot plots display the average expression of top gene markers that characterize each sub-cluster. (C) Branching trajectory analyses using Monocle3 exposed a genotype-dependent bifurcation within the oligodendrocyte lineage, diverging at subcluster 5 into distinct trajectories for WT and dKO cortices. (D) Relative cell proportions within each subcluster. (E) Dot plots highlight key markers implicated in oligodendrogenesis, cellular communication, and myelination processes across the subclusters. Genes pinpointed by the branching analysis are visualized through (F) violin plots, comparing expression levels in WT and dKO oligodendrocyte clusters. (G) They are further visualized with feature plots, displaying the log normalized data scaled to the percent of maximum expression. Average expression in (B) and (E) calculated as log(mean + 0.1). Values represent means ± SD. * p < 0.05 and *** p < 0.001.

Last, to investigate additional genomic links between these genes and AHDS phenotypes, we analyzed single nucleotide polymorphism (SNP) associations in the NHGRI-EBI genome-wide association studies (GWAS) catalogue [37]. From 670 traits, we selected 18 relevant to AHDS, including neurological symptoms and brain morphology [5]. *Dcc, Cdh13, and Magi2* appeared in over 10 traits, primarily psychiatric related (Suppl. Fig. 8A, Supplementary File 6). We further used the human genetic evidence score to predict the gene’s implications on a specific trait [38]. *Cdh13*, *Magi2*, and *Cntn2* showed evidence for hypothyroidism or thyroid traits. *Dcc* was strongly related to cognitive and psychiatric traits (Suppl. Fig. 8B, Supplementary File 7). In addition, using the polygenic priority score to demonstrate which genes compile more human evidence at a functional level [39], we found high scores for psychiatric disorders for *Magi2* and *Dcc* (Suppl. Fig. 8C, Supplementary File 8).

## Discussion

In AHDS and its murine models, developmental TH deficiency has severe effects on brain morphology and function [12][14]. By applying single-nucleus RNA sequencing to WT and MCT8 and OATP1C1 dKO mouse cortices at P21, we aimed to specifically unravel AHDS disease mechanisms at cellular resolution during a critical stage when disruptions in neurogenesis and myelination are already evident yet not confounded by aging effects [19][40]. Our snRNAseq analysis revealed key molecular perturbations in the AHDS cortex, including shifts in GABAergic and glutamatergic cell proportions and bifurcation within the oligodendrocyte lineage, resulting in fewer mature oligodendrocytes with deregulated genes related to the ECM and associated with brain dysfunctions. Additionally, aberrant primary cilia formation and an imbalance in GABAergic versus glutamatergic signaling suggesting altered cell-cell communication, underscore the critical role of thyroid hormone signaling in maintaining network homeostasis in the cerebral cortex and cerebral nuclei.

Such effects were likely not only cell-autonomous, judged by the discrepant T3 transporter expression profiles. In the adult mouse brain, MCT8 protein expression is largely restricted to blood vessels, tanycytes, and the choroid plexus [41][42]. Accordingly, how TH is transported from the BBB toward distant brain areas and cell types continues to remain unknown. At the BBB, T3 and T4 are predominantly transported by MCT8 in ependymal and endothelial cells and tanycytes, while astrocytes and endothelial cells expressing OATP1C1 contribute to the transport of T4 and local conversion to T3. In the context of the mouse model of AHDS, where the primary pathways for T4 entry via OATP1C1 and T3/T4 uptake via MCT8 are deficient, severely diminished T3 and T4 levels in the brain parenchyma align with a widespread disturbance in T3 signaling throughout the cortex and striatum, verified by our comparison to existing bulk sequencing data on mouse cortex and striatum of another MCT8/OATP1C1 dKO mouse model, and mice with systemic hypothyroidism. Processes of brain development, synaptic signaling, as well as cell-adhesion were deregulated throughout the neuron types. However, we also saw diverging gene expression profiles between SH and the dKO models. Future studies, preferably at single-nucleus resolution, are thus warranted to untangle the disturbances caused by SH alone and by MCT8 and OATP1C1deficiency. These may focus on the role of genes implicated in cilia formation and function, which were perturbed in MCT8/ OATP1C1 dKO but not SH. In the dKO cortex, we observed increased cilia numbers, and in the somatosensory cortex longer ciliary length. Primary cilia, cellular antennae involved in brain development via e.g., Wnt and Shh pathways, are linked to neurodevelopmental disorders when dysfunctional [43][44]. Roles for cilia length are proposed, longer cilia may enhance signaling by increasing receptor surface area, while shorter cilia may concentrate ligands [45]. To date, structural and functional abnormalities in primary cilia have not been associated with the neurological deficits in the AHDS. The observed alterations may either represent cellular compensatory mechanisms, or functional drivers for the AHDS disease phenotype.

Motor and cognitive impairments due to TH deprivation are often linked to functional deficiencies in GABAergic neurons, most notably parvalbumin expressing neurons [10][19][46]. Pvalb protein staining was shown to be diminished in P12 [19] and here in P21 dKO cortices. Of note, at P21 gene expression levels for both *Slc16a2* and *Slco1c1* were near absent in Pvalb neurons, indicating that MCT8/OATP1C1-deficiency primarily influences Pvalb neuron maturation before P21 or that Pvalb neurons are more susceptible to the generally hypothyroid state in the brains of MCT8/OATP1C1 dKO mice.

In contrast to Pvalb neurons, proportions of STR D1 and D2 neurons were increased in dKO mice. These clusters are projection neurons originating from striatal tissue that remained attached to the cortex after dissection [47]. Unlike Pvalb neurons, STR D1 and D2 neurons derive from the lateral ganglionic eminence (LGE), with earlier onset of neurogenesis [48]. The observed changes in cell proportion of these neurons might indicate that the time-point during development is crucial for the extent of MCT8/OATP1C1 mediated effects on neuron maturation. The role of TH during LGE progenitor formation, and the mechanisms underlying the increased neuron numbers in the hypothyroid dKO brain and their implications for the AHDS disease phenotypes remain unclear, warranting further investigation.

GABAergic input was inferred to be increased for most cell groups in cortex and striatum and the output was increased for CNU GABA and LSX GABA. Combined with the increased cell proportion of STR D1 and D2 neurons, such a disbalance in GABAergic vs glutamatergic signaling is likely contributing to movement deficits, as striatum-residing dopamine-expressing spiny neurons are part of the basal ganglia system responsible for movement control [49].

While less studied in MCT8-deficiency, hypothyroidism is known to impair also glutamatergic neuron generation and migration [8][50][51]. These disruptions result in structural anomalies in cortical layers, likely contributing to the cognitive and motor impairments observed in AHDS patients. In dKO mice, cortical layers I-IV are thinner already at P12 [12]. We suspected the reduced number of glutamatergic L4/5 IT CTX neurons to be a potential reason but could not confirm decreases in P21 brain slices. Glutamatergic neurons exhibited the highest number of DEGs, and cell-cell communication analyses revealed reduced signaling, inferred from decreased expression of glutamate-synthesizing enzymes, vesicular transporters, and receptors. Overall, our data highlight the perturbations of cell-cell communication patterns by glutamatergic neurons. They are consistent with a recent study on the adult mouse motor cortex that linked T3 treatment with altered glutamatergic gene expression patterns and increased excitatory postsynaptic currents, while a mutation reduced this effect [27].

Alongside the shift from excitatory to inhibitory signaling, we observed changes in synapse formation and plasticity, inferred from neurexin-neuroligin interactions. These synaptic adhesion molecules regulate synaptogenesis and transmission, and their dysfunction is linked to autism, schizophrenia, and seizures [52][53]. Downregulated interactions suggest impaired connectivity and signal transduction in the P21 dKO cortex, adding to the changes in neuron number and signaling balance as a potentially explanation for the motor and behavioral deficits. Elevated neurexin or neuroligin expression can also cause neurological impairments. Notably, Nrx2, increased for many dKO clusters, acts as an inhibitory neurexin in the hippocampus, its overexpression leading to seizures [53].

Myelination is a key process in nervous system development and is often reduced in the AHDS. The reduction in myelinated axons in dKO mice was reported to result from impaired oligodendrocyte maturation, with an increased OPC pool and fewer mature oligodendrocytes [12][54]. Next to corroborating this, we uncovered a bifurcation in the oligodendrocyte lineage, forming two mature populations distinct between WT and dKO. Notably, however, many commonly studied myelination-related genes were normally expressed, suggesting a largely intact function. Reduced MBP staining thus appears to result from fewer oligodendrocytes rather than impaired myelination potential, consistent with published reports showing longer myelin sheaths but fewer myelinated axons [12]. The distinct gene expression patterns for the WT and dKO oligodendrocyte lineage may provide a framework for deeper analyses. Polygenic priority scores of the GWAS analysis predict axon-guidance receptor *Dcc* as potential candidate gene to explain some of the bifurcation and myelination phenotype observed in the dKO and AHDS, alongside *Magi2, Cdh13*, *Cntn2* as well as *Lama2* and *Litaf.* Additional candidate genes include the *Pisd-ps3* pseudogene to the phosphatidylserine decarboxylase *Pisd*, which is upregulated as top DEG in multiple dKO clusters*. Pisd* is involved in lipid transport and metabolism [55][56], two critical processes necessary for the extensive lipid production during myelination [57]. Notably, although the role of the *Pisd* pseudogene remains elusive, *Pisd-ps3* and *Pisd* are inversely expressed in some dKO organs. *Pisd-ps3* has been identified in other datasets, showing upregulation in the dKO mouse cortex and striatum [24] as well as in the hypothalamus of *THRa1* mutant mice [58], highlighting a potential regulatory relationship with TH signaling. Interestingly, T3-treatment in the adult motor cortex also led to upregulation [27]. Further exploration of *Pisd-ps3* is warranted to determine whether this *Pisd* pseudogene or *Pisd* itself exert a functional role in lipid metabolism and myelination, potentially under TH control.

### Conclusion

This study provides a valuable resource and significant insights into the neurodevelopmental processes influenced by the combined deletion of MCT8 and OATP1C1 in our AHDS mouse model. Although these two transporters were not expressed in every cluster, the majority of cell types exhibited numerous DEGs and showed impairments in neurodevelopmental processes associated with TH deficiency. We validated multiple genes perturbed by the concomitant MCT8 and OATP1C1 deletion by using public domain bulk sequencing data and showed a significant overlap, but also clear distinctions to those genes deregulated by systemic hypothyroidism. Overall, the single-cell resolution provided unprecedented insight into the specific cell types most affected and the nature of their disruptions. We could expand previously known disease aspects by revealing a bifurcation in the oligodendrocyte lineage, uncovered changes in the number and connectivity of GABAergic but also glutamatergic neurons, adding to the well-described GABAergic Pvalb neuron impairments, and revealed a previously unknown primary cilia dysfunction. Collectively, such perturbations in the formation of neuronal networks, oligodendrocyte differentiation and myelination, and cilia-driven pathways essential for brain development and neuronal communication are likely contributors linking impaired thyroid hormone signaling to the neurological deficits in the AHDS.

### Limitations

Sequencing the entire cortical hemisphere allowed us to analyze a broad range of cell types but resulted in lower granularities in respect to defined cortical areas. Additionally, translating these findings to humans remains challenging due to differences in MCT8 and, particularly, OATP1C1 expression patterns, as well as the greater complexity of the human cortex compared to the mouse cortex. Comparisons to human GWAS provide valuable insights, but further experimental manipulation of these genes is essential to confirm their roles in AHDS.

## Material and Methods

### MCT8/OATP1C1 dKO animal model

Global MCT8/OATP1C1 dKO mice represent a mechanistically faithful model of AHDS that enable detailed analyses of disease perturbations at cellular resolution, as previously reported [14]. MCT8 KO and OATP1C1 KO mice were crossed to generate the dKO, and regularly backcrossed to a pure C57BL/6J background. Genotyping primers are available in Table 1. All mice were group-housed on a 12:12-h light-dark cycle at 23°C and fed standard chow diet *ad libitum*. At the age of 21 days, immediately after cervical dislocation, blood was taken in EDTA-coated syringes and centrifuged at 2,000 x g for 10 min at 4°C to separate the plasma. The brain was extracted and weighed. The cortex and all other brain areas were thoroughly dissected using anatomical references for guidance, weighed and then snap frozen in liquid nitrogen and stored at −80°C. Organs were dissected and frozen and stored the same way. All procedures involving animal handling were approved by the committee for Care and Use of Laboratory Animals of the Government of Upper Bavaria, Germany.

**Table 1.**
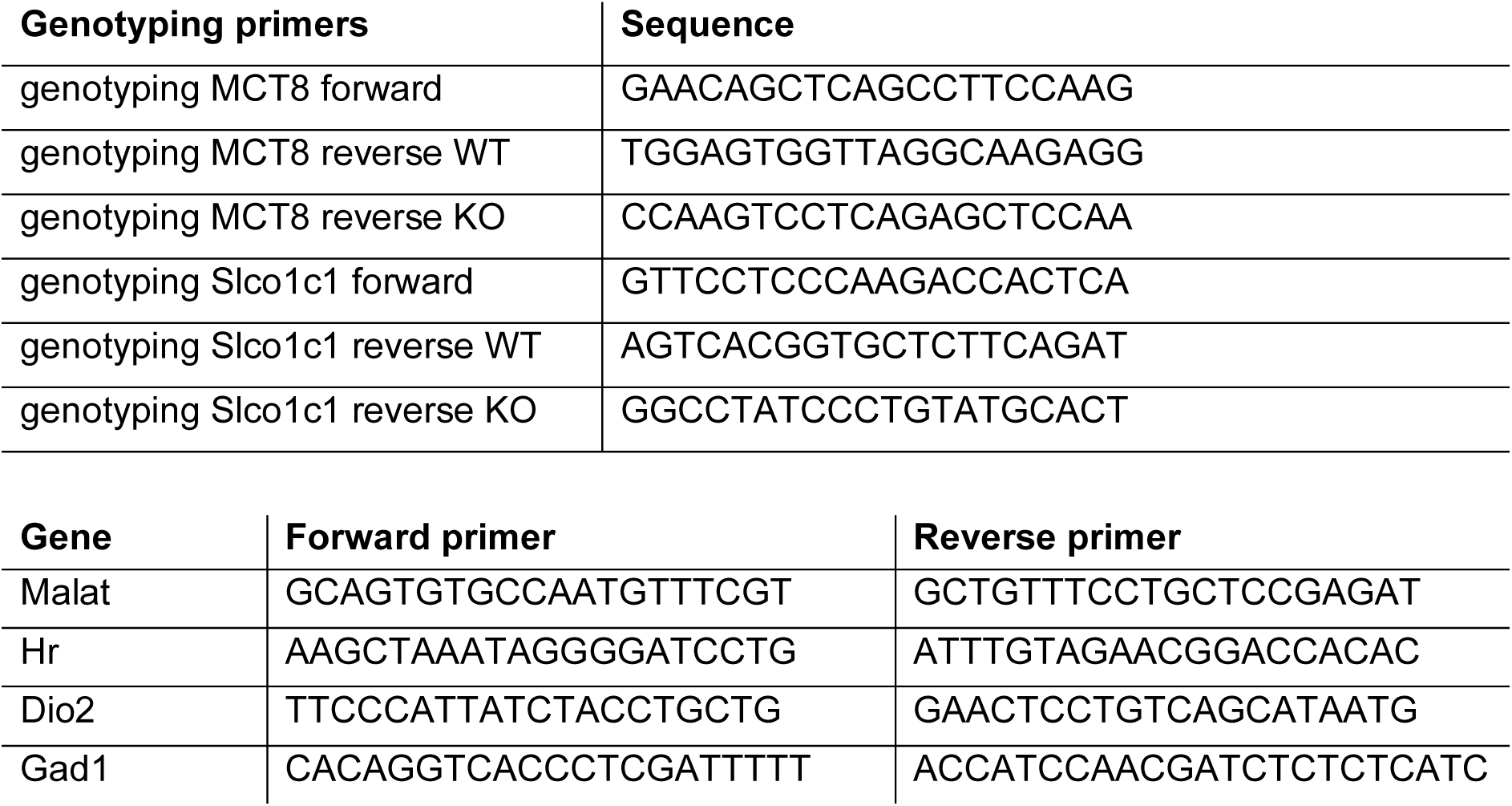
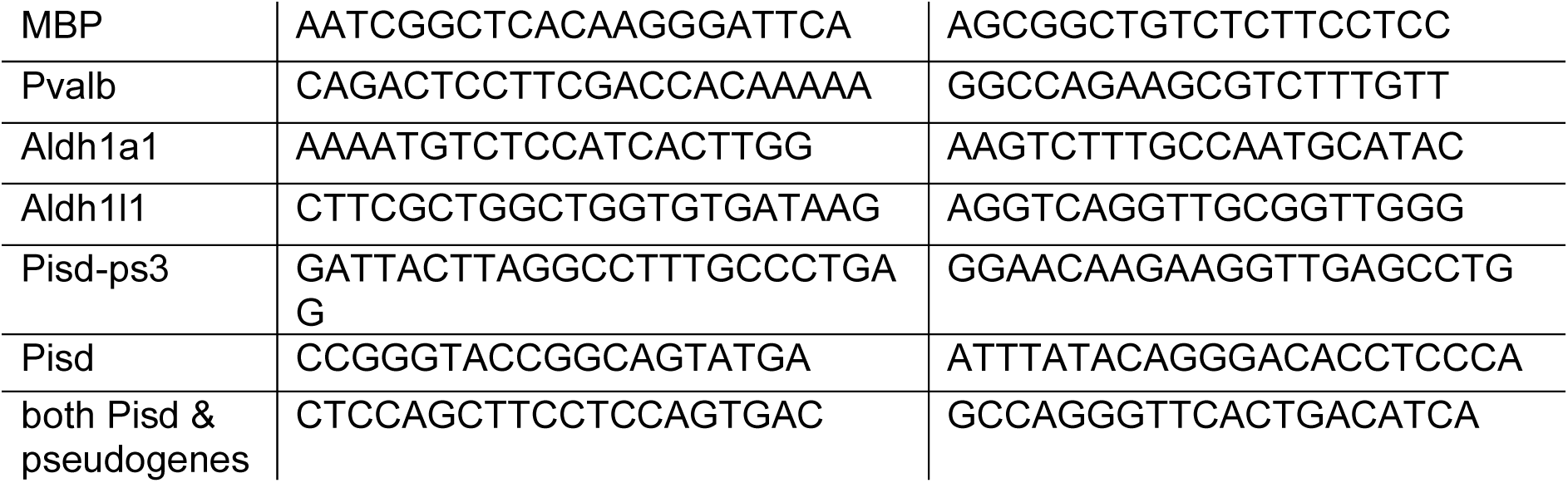
Primers.

### tT4 and tT3 measurement by LC-MS/MS

Flash-frozen brain areas were homogenized to powder at −200°C, of which ∼100 mg was used for the extraction of TH using standard solutions and clean-up procedures as previously described [59]. For plasma analysis, 50 µL of mouse plasma were mixed with 60 µL of internal standard (10 pg/µL) and an antioxidant solution (0.15 mL; 25 mg ascorbic acid + 25 mg citric acid + 25 mg dithiothreitol in 1 mL H_2_O). The mixture was vortexed for 10 sec and equilibrated for 1h at 4°C. Subsequently, 25 µL of ZnCl_2_ (2M in water) and 200 µL of methanol were added and additionally incubated for 30 min at 4°C. Then, the sample was centrifuged at 3000x g for 10 min and supernatant collected. The solid residue was further resuspended in a solution of methanol:water ((1:1), 0.2 mL), centrifuged, and then extracted as described before. Chloroform (0.6 mL) was added to the combined extracts and the mixture was centrifuged again (3000 x g, 10 min). The upper level was decanted while the lower phase was re-extracted (methanol:water ((1:1), 0.2 mL). The pooled upper phases were diluted with 1.5 mL pure water followed by the addition of phosphoric acid to achieve a final concentration of 2%. After vortexing, the mixture was loaded onto a Bond Elut Plexa PCX cartridge, which was preconditioned sequentially with 2 mL of pure methanol and 2 mL of water. The cartridge was first washed with 2 mL of 2% formic acid in water and then with 2 mL of methanol:acetonitrile (1:1, v/v). Analytes were eluted into a vial with 1 mL of 5% ammonium hydroxide in methanol:acetonitrile (1:1, v/v). The solvent was evaporated and the compounds re-dissolved in 60 µL of a mixture of 20% acetonitrile in water.

The detection and quantification of seven THs: L-Thyroxine (T4), 3,3′,5-triiodothyronine (T3), reverse 3,3’,5’-triiodothyronine (rT3), 3,3’-diiodo-L-thyronine (T2), 3,5-diiodo-L-thyronine (rT2), 3-iodo-L-thyronine (T1) and 3-iodothyronamine (T1AM) was performed with a Sciex QTrap 5500 mass spectrometer interfaced with an Agilent 1290 Infinity II LC system or a Shimadzu Nexera X2 LC system. For both configurations, the flow rate was set at 0.3 mL/min, the column used was a Zorbax Eclipse Plus C18 (2.1×50 mm, 1.8 uM) with a column temperature fixed at 40°C. Injection volumes were 10 μL. The mobile phases were water (A) and acetonitrile (B) each containing 0.1% formic acid (v/v). Gradient elution was the same for both instrumentations as reported in Table 2. Analytes were detected using an electrospray ionization source (ESI) in positive mode. The tandem mass spectrometer was operated under multiple reaction monitoring mode (MRM). The MS/MS parameters for both configurations were the same and indicated in Table 3. Data acquisition, linearity of the standard curves and quantification of the samples were performed using Analyst software.

**Table 2.** Elution parameters for the analysis of THs.

| TIME (min) | PUMB B (%) |
| --- | --- |
| 0.1 | 5 |
| 3 | 5 |
| 4 | 30 |
| 5.5 | 35 |
| 6.5 | 35 |
| 7.0 | 40 |
| 8.0 | 100 |
| 10.0 | 100 |
| 10.2 | 5 |
| 13.2 | 5 |

**Table 3.** Optimized MS/MS parameters for THs. For each compound, ion-transitions are shown as m/z for the parent ion and two product ions (for quantification (q) and confirmation (c)). Compound optimized values for retention time (t_R_), declustering potential (DP), collision energy (CE), and collision cell exit potential (CXP).

| Compound | $t_R$<br>(min)<sup>a</sup> | Parent<br>ion (m/z) | Product<br>ions (m/z) | DP<br>(V) | CE<br>(V) | CXP<br>(V) |
| --- | --- | --- | --- | --- | --- | --- |
| Target compounds |  |  |  |  |  |  |
| T<sub>4</sub> | 6.90 | 778 | 732(q) | 75 | 40 | 20 |
|  |  |  | 634 (c) | 75 | 40 | 20 |
| T<sub>3</sub> | 6.35 | 652 | 606 (q) | 70 | 35 | 15 |
|  |  |  | 508 (c) | 70 | 35 | 15 |
| rT<sub>3</sub> | 6.07 | 652 | 606 (q) | 70 | 35 | 15 |
|  |  |  | 508 (c) | 70 | 35 | 15 |
| 3,3'-T<sub>2</sub> | 5.67 | 526 | 480 (q) | 65 | 40 | 20 |
|  |  |  | 382 (c) | 65 | 40 | 22 |
| 3,5-T<sub>2</sub> | 5.34 | 526 | 480 (q) | 65 | 35 | 20 |
|  |  |  | 382 (c) | 65 | 35 | 22 |
| T<sub>1</sub> | 5.11 | 400 | 256 (q) | 70 | 35 | 20 |
|  |  |  | 354 (c) | 70 | 35 | 20 |
| T<sub>1</sub>AM | 5.22 | 356 | 339 (q) | 75 | 40 | 35 |
|  |  |  | 212 (c) | 75 | 40 | 25 |
| Internal standards |  |  |  |  |  |  |
| ML-T<sub>4</sub> | 6.90 | 784 | 738 | 75 | 35 | 20 |
| ML-T<sub>3</sub> | 6.35 | 658 | 612 | 70 | 35 | 15 |
| ML-rT<sub>3</sub> | 6.07 | 658 | 612 | 70 | 35 | 15 |
| ML-3,3'-T<sub>2</sub> | 5.67 | 532 | 486 | 65 | 40 | 25 |
| ML-T<sub>1</sub>AM | 5.22 | 362 | 345 | 75 | 40 | 35 |

<sup>a</sup> Retention time for the Shimadzu Nexera X2 LC.

### RNA isolation and gene expression analysis

Tissue samples were pulverized on liquid N_2_, ∼ 10 mg was weighed and further homogenized in a Tissue Lyser II for 3 min at 30/sec with 500 µl Qiazol. After a 5 min incubation step at RT, 100 µl chloroform was added to each sample, vigorously mixed, and incubated for another 3 min at RT. After resolving the phases by centrifugation at 12000 x g for 15 min at 4°C, the supernatant was used for RNA isolation with the NucleoSpin RNA isolation kit (740955, Machery-Nagel) according to the manufacturer’s instructions. Subsequently, RNA concentrations were measured on a Nanodrop and the same amount of RNA for each sample was reverse-transcribed into cDNA using QuantiTect® Reverse Transcription Kit (205311, QIAGEN). Gene expression was quantified using the SYBR® Green PCR Master Mix (Applied Biosystems™) according to the manufacturers guidelines with a reduced reaction volume of 5 µl and target specific primers (Table 1) in a QuantStudio 7 Flex Real Time PCR System (Applied Biosystems™). Differential gene expression was calculated using the 2-ΔΔCt method normalized to *Malat1*.

### Immunofluorescent staining and RNAscope™ in cortex slices

Mice were euthanized with an overdose of ketamine/xylazine and perfused through the heart using a peristaltic pump, first with ice-cold PBS, then with 4% paraformaldehyde (PFA). Brains were post-fixed overnight in 4% PFA at 4°C, for immunofluorescent staining this was followed by equilibration with 30% sucrose in Tris-buffered saline (TBS, pH 7.2) for 48 h before sectioning into 30 μm coronal slices using a cryostat (CM3050S; Leica, Germany). Anterior brain sections (Bregma 0.86 to 0.62) and for cilia analysis also middle areas (Bregma −1.58 to −2.06) were selected and washed with TBS, then mounted on superfrost slides and dried shortly. Antigen retrieval was performed by boiling the slides in 10 mM sodium citrate buffer pH 6 in the microwave for 10 min. After cooling and shortly rinsing with TBS the tissue was blocked and permeabilized using a solution of 0.25 % (w/v) porcine gelatin and 0.5 % (v/v) TritonX 100 in TBS for 1-2 h at room temperature. For Rorb staining, mouse-on-mouse blocking was performed using the blocking solution by Invitrogen (R37621) according to the manufacturer’s instructions. For free-floating staining (MBP, Olig2, Iba1), sections were washed with TBS, then permeabilized and blocked as well. Subsequently, slices and slides were incubated overnight at 4°C with primary antibodies (see Table 4 in the solution containing 0.25% porcine gelatine and 0.5% Triton X-100 in TBS. The following day, sections were rinsed three times in TBS and incubated with respective secondary antibodies and DAPI (see Table 4) for 2 h. Sections were washed three times in TBS, free-floating sections were mounted, and the slides were covered in Elvanol mounting medium (150 mM Tris, 12% Mowiol 4-88, 2 % DABCO) and sealed under a coverslip.

**Table 4.**
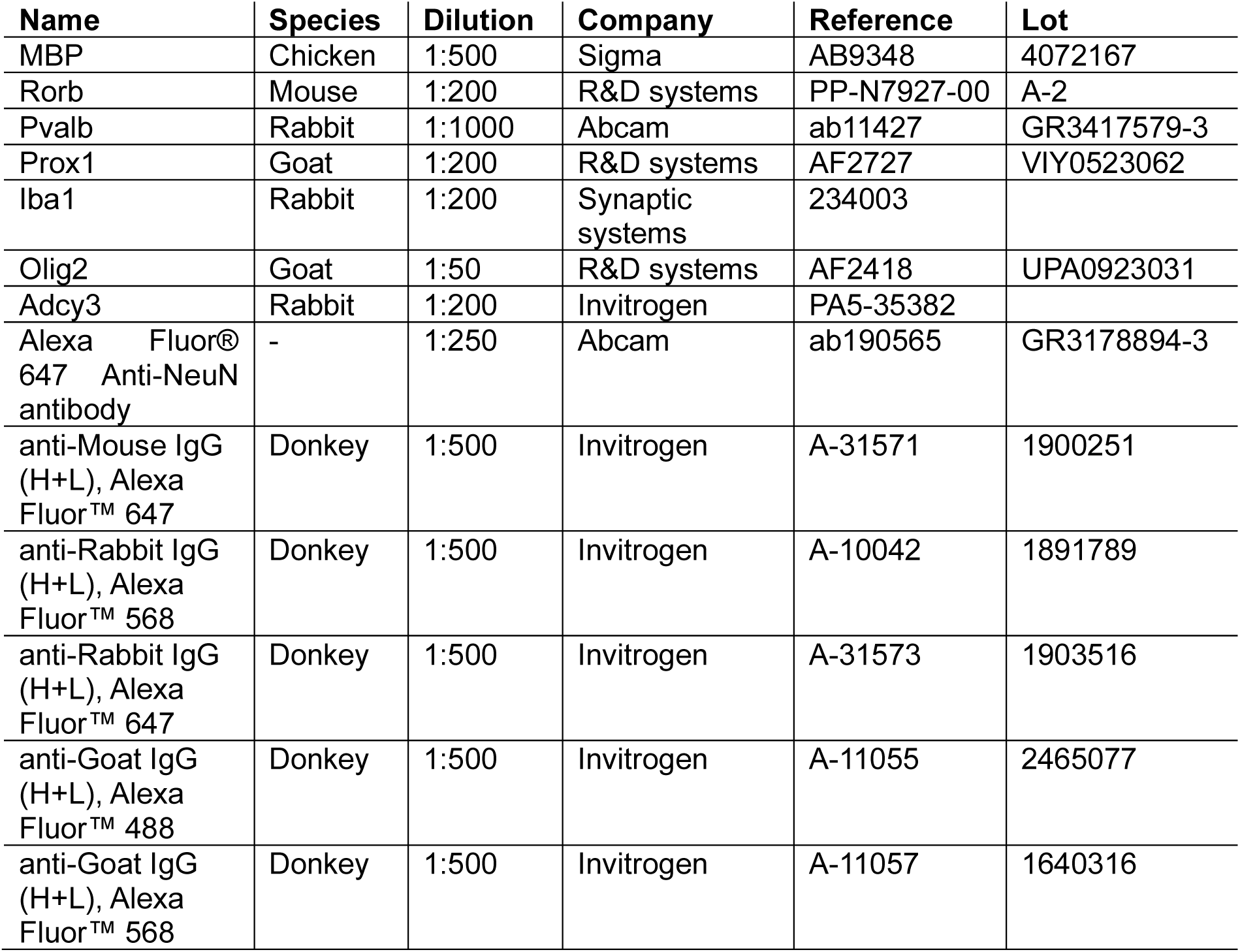
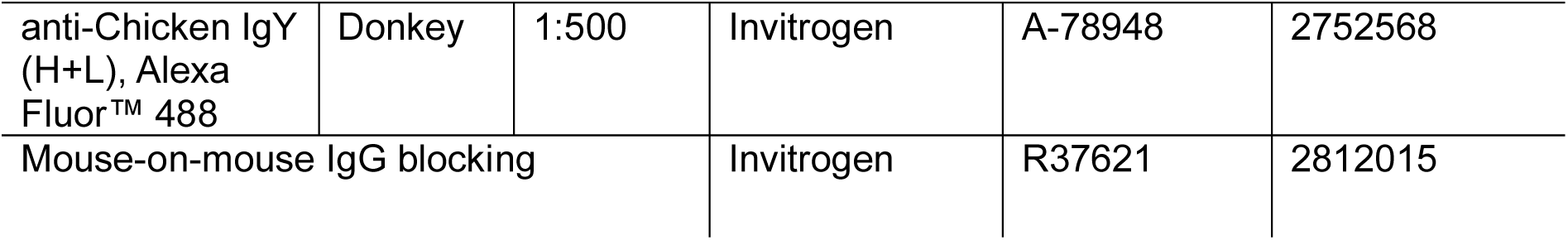
Antibodies.

| **Name** | **Species** | **Dilution** | **Company** | **Reference** | **Lot** |
| --- | --- | --- | --- | --- | --- |
| MBP | Chicken | 1:500 | Sigma | AB9348 | 4072167 |
| Rorb | Mouse | 1:200 | R&D systems | PP-N7927-00 | A-2 |
| Pvalb | Rabbit | 1:1000 | Abcam | ab11427 | GR3417579-3 |
| Prox1 | Goat | 1:200 | R&D systems | AF2727 | VIY0523062 |
| Iba1 | Rabbit | 1:200 | Synaptic systems | 234003 |  |
| Olig2 | Goat | 1:50 | R&D systems | AF2418 | UPA0923031 |
| Adcy3 | Rabbit | 1:200 | Invitrogen | PA5-35382 |  |
| Alexa Fluor® 647 Anti-NeuN antibody | - | 1:250 | Abcam | ab190565 | GR3178894-3 |
| anti-Mouse IgG (H+L), Alexa Fluor™ 647 | Donkey | 1:500 | Invitrogen | A-31571 | 1900251 |
| anti-Rabbit IgG (H+L), Alexa Fluor™ 568 | Donkey | 1:500 | Invitrogen | A-10042 | 1891789 |
| anti-Rabbit IgG (H+L), Alexa Fluor™ 647 | Donkey | 1:500 | Invitrogen | A-31573 | 1903516 |
| anti-Goat IgG (H+L), Alexa Fluor™ 488 | Donkey | 1:500 | Invitrogen | A-11055 | 2465077 |
| anti-Goat IgG (H+L), Alexa Fluor™ 568 | Donkey | 1:500 | Invitrogen | A-11057 | 1640316 |
| anti-Chicken IgY (H+L), Alexa Fluor™ 488 | Donkey | 1:500 | Invitrogen | A-78948 | 2752568 |
| Mouse-on-mouse IgG blocking |  |  | Invitrogen | R37621 | 2812015 |

For RNAscope™, post-fixed brains were equilibrated step-wise in 10%, 20%, and 30% sucrose, before sectioning into 10 µm coronal slices using a cryostat. Slices were immediately mounted on slides and air dried at −20°C. RNAscope™ was performed according to the user manual of the RNAscope™ Multiplex Fluorescent Kit v2 (# 323100) and 4-Plex Ancillary Kit (#323120) from ACD. The probes and Opals™ used are listed in table 5. For combined staining with the NeuN antibody, the protease-free workflow of PretreatPro™ was used instead of protease III treatment. Acquisition was performed on a Leica SP8 or Zeiss Axio Scan 7 slidescanner.

**Table 5.** RNAscope™ probes and Opals™.

| Probe Name | Channel | Catalog # | Lot Number | Opal™ used |
| --- | --- | --- | --- | --- |
| Mm-Rorb | C1 | 444271 | 24275A | Opal 520 |
| Mm-Litaf | C1 | 1119771-C1 | 24285B | Opal 520 |
| Mm-Penk | C2 | 318761-C2 | 24275B | Opal 570 |
| Mm-Mobp | C2 | 431721-C2 | 24275C | Opal 570 |
| Mm-Pdyn | C3 | 318771-C3 | 24274C | Opal 780 |
| Mm-Lama2 | C3 | 424661-C3 | 24285A | Opal 690 |
| Opals™ (Akoya Biosciences) |  | Catalog # | Lot Number |  |
| Opal 520 Reagent Pack |  | FP1487001KT | 230919042 |  |
| Opal 570 Reagent Pack |  | FP1488001KT | 230816015 |  |
| Opal 690 Reagent Pack |  | FP1497001KT | 230918002 |  |
| Opal 780 Reagent Pack |  | FP1501001KT | 240307016 |  |

### Quantification of immunofluorescent staining, RNAscope™, and cilia detection

For the quantification of stained cells, we used QuPath (v0.5.1) [60] with automatic cell detection or trained classifiers. An annotation was made according to the Allen Mouse Brain Reference Atlas [61] within the motor cortex, with the size adapted to the thinner cortex in dKO mice. Positive cells were then detected using “Cell detection” in the respective channels in case of stainings with defined intensity ranges. For Rorb-staining we had to use the “Train object classifier” function of QuPath, since there were many different intensities for positive cells. The counts for positive cells were then normalized to the annotation area in WT and dKO. For Iba1 detection we used Fiji [62] and first prepared the image with a rolling ball background subtraction (radius 15, disable smoothing), followed by automatic thresholding to define the Iba1 stained ROI. This ROI was applied to the original image to measure the integrated density of the stained microglia.

To quantify RNA levels after RNAscope™, QuPaths’ subcellular detection was used to get estimated number of spots which were normalized to cell numbers in the detection area using DAPI detection or to positive cells using the Mobp probe or NeuN staining.

To detect and analyze cilia we used the CiliaQ plugin for Fiji [63]. The Z-stack images with 1 µm distance were prepared using CiliaQ Preparator v0.1.2, with “Gaussian blur - radius” of 2.8 and “Segmentation method Triangle”. Cilia detection (CiliaQ v0.1.7) was done with a minimum size of 6 voxels, increasing the range for connecting cilia, excluding cilia touching x or y or z borders, and pre-set skeleton-based results parameters (gauss filter XY = 2, Z sigma = 0).

### Fluorescence-Activated Nuclei Sorting (FANS) of cortical nuclei for single nucleus RNA sequencing

For nuclear isolation, 9 WT and 9 dKO frozen cortical hemispheres were each transferred to a Dounce homogenizer containing 3 mL of freshly prepared ice-cold nuclei isolation buffer (0.25 M sucrose, 25 mM KCl, 5 mM MgCl2, 20 mM Tris pH 8.0, 0.4% IGEPAL 630, 1 mM DTT, 0.15 mM spermine, 0.5 mM spermidine, 1x phosphatase & protease inhibitor tablet, 0.4 units RNasin Plus RNase Inhibitor, 0.2 units SuperAsin RNase inhibitor) [64]. Homogenization was achieved by 5 pestle strokes with a looser pestle, followed by an incubation step on ice for 5 min and 20 more strokes with a tighter pestle. The homogenates were filtered through a 20 µm cell strainer and centrifuged at 1000 x g for 10 min at 4°C. The nuclear pellet was resuspended in 500 µl of staining buffer (RNAse-free PBS pH 7.4, 0.15 mM spermine, 0.5 mM spermidine, 0.4 units RNasin Plus RNase Inhibitor, 1.5% RNAse-free BSA, 1 µg/µL DAPI). 150 µl nuclei suspension of three samples was pooled and nucleus integrity was assessed in the DAPI channel under a Zeiss microscope (Axio Scope, Zeiss, Germany). A total of three WT and three dKO samples were sorted in a FACS-Aria III (BD Biosciences) using a 70 µm nozzle. Appropriate gating for single nuclei was set using side scatter height and area parameters, as well as DAPI signal for doublet discrimination. Nuclei were collected on a 10 µl cushion of BSA and Protector RNAse inhibitor, reaching a final concentration of <1% BSA and 0.2 units Protector RNAse inhibitor. Samples were then stored on ice.

Immediately after sorting, GEM formation and library preparation was performed using the Chromium Next GEM Automated Single Cell 3′ Reagent Kit v3.1 (Dual Index, 10× Genomics, #1000268) according to the manufacturer’s instructions. The resulting libraries underwent 150-bp paired-end sequencing of read 2 on a NovaSeq 6000 (Illumina).

### Preprocessing of single-nucleus transcriptomics data

Gene-barcode count matrices were generated by mapping reads to the mouse reference genome GRCm39 (Ensembl release 109) using the CellRanger (v.6.1.2) following sequencing. The downstream analysis was performed using Seurat version 4 [65]. Nuclei having fewer than 1000 and more than 6000 genes, more than 30,000 UMIs, and more than 5% mitochondrial reads were filtered out. Doublet Finder V2.0 [66] was employed to remove doublets, and data was normalized using SCTransform [67]. The top 3000 highly variable features were selected for integration using canonical correlation analysis (CCA). Data integration was done using IntegrateData. First 50 principal components were used to identify neighbors and clustering was done using louvain algorithm. The data was visualized using Uniform Manifold Approximation and Reduction (UMAP).

### ClusterMarkers and Differential expression analysis

Cluster markers were determined using the Wilcoxon rank sum test with the FindAllMarkers function. Log2FC threshold of 0.25, and a detection in at least 25% of the nuclei was set as cut-off. For multiple testing, Bonferroni correction was used. Cell type annotation was done using reported markers in literature. Differential expression analysis was performed using DESeq2 [68] with default settings. Genes having abs(Log2FC) >= 1 and adjusted p-value less than 0.05 were considered as differentially expressed between conditions. Volcano plots for DEGs were generated using R package EnhanceVolcano [69]. R packages ggplot2, tidyverse, and dplyr were used for data visualization and wrangling [70][71][72].

### Cell proportion

Cell type proportions were calculated using propeller function of Bioconductor package speckle [73]. P-values were calculated using t-test.

### Gene set enrichment analysis

To identify dysregulated pathways and processes, gene set enrichment was performed using GSEA v4.3 [74][75]. Signal2Noise Metric was used to generate ranked list. This ranked list was used against Hallmark pathways, GO biological processes, KEGG, BioCarta, Reactome datasets from MsigDB for enrichment [76]. Gene sets having p-value < 0.05 and FDR < 0.25 were considered as significantly enriched.

### Correlation of pseudobulk data to MO dKO and SH dataset

After preprocessing, filtering, and integration of the WT and dKO cortical nuclei, we aggregated counts at the sample level by summing raw UMI counts across all nuclei belonging to predefined anatomical groupings. Nuclei assigned to cortical excitatory and inhibitory clusters (CTX) together with non-neuronal cell types (NN) were combined to create a pseudobulk cortex dataset, whereas nuclei annotated as cerebral nuclei (CNU) formed the pseudobulk striatum dataset. For each sample, gene-level counts were summed and subsequently normalized using the DESeq2 median of ratio to approximate bulk RNA-seq– like expression distributions. Differential expression analysis between WT and dKO pseudobulk samples was performed using DESeq2, treating each biological replicate as an independent observation. We compared our pseudobulk cortex and striatum signatures to the publicly available P21 bulk RNA-seq datasets from Morte et al. [24]. Log2 fold changes from our pseudobulk analyses were aligned with those reported by Morte et al. and overlap statistics and directional correlations were computed based on the intersecting sets of differentially expressed genes.

### Neuron-neuron communications

Neuron-neuron communication was inferred using NeuronChat [31]. A communication strength matrix for all ligand-target pairs was generated with run_NeuronChat, employing 100 permutation tests and controlling the FDR at 0.05. Overall communication strength was evaluated by interaction counts and weights using compareInteractions_Neuron. Individual pairwise interactions were identified with rankNet_Neuron. Shared and specific interactions were assessed using computeNetSimilarityPairwise_Neuron, and manifold learning of signaling networks of interaction pairs was performed with netEmbedding. K-mean clustering was employed to cluster interaction pairs using netClustering, and visualized with netVisual_embeddingPairwise_Neuron.

### Trajectory analysis

To better understand the differences in Oligo lineage between WT and dKO, trajectory analysis was performed using Monocle3 [77][78]. Nuclei annotated as oligodendrocyte precursor cells, newly formed oligodendrocytes, and oligodendrocytes were extracted and a cell data set object was generated. Dimension reduction was done using first 16 principal components and align_cds() was used to align nuclei. Nuclei were clustered cells at a resolution of 0.0007 and trajectory was learned using learn_graph(). Data was subsetted to clusters 1, 2, 3, and 5 for trajectory branch point analysis. Genes that showed difference in expression in the chosen trajectory area were calculated by graph_test().

### Association with human traits using GWAS

For the genes differentially expressed in the WT and oligodendrocyte subclusters, we looked their association with reported phenotypes using genome-wide association studies. The NHGRI-EBI GWAS Catalog contains variants-trait associations from more than 5000 human traits and over 45000 GWAS studies [37]. Summary statistics (Supplementary File 6) were downloaded from the NHGRI-EBI GWAS Catalog on 06/10/2024. Traits relevant for AHDS were filtered and summarized manually.

We then used the human genetic evidence score, which analyzes, by Bayes factor, the common and rare variants associated with a trait collected from 758 phenotypes, 654 genetic datasets, and 7745 genomic datasets [38]. The results are filtered by AHDS relevant traits and summarized by their evidence score.

Further refinement was done using the polygenic priority score, which is a computation method that identifies GWAS variants/locus and analyses them with regulatory evidence (QTLs and tissue-specific chromatin conformation) with model organism phenotypes, drug interactions, among other data [39]. The resulting scores were plotted for traits relevant for AHDS.

## Author Contributions

AM, EP, SCS & PTP conceptualized and designed all experiments; EP performed bioinformatical analyses, with support by BO and DL; AM, SCS, CM conducted murine studies and stainings; NM, MB performed nuclear isolations and library preparations for snRNAseq, as well as RNAscope™ experiments; MDeA conducted LC-MS based TH analyses; RGA assessed GWAS associations; GM, PK, RO and TDM provided experimental guidance and scientific advice, and revised the article; AM, EP, SCS and PTP wrote the manuscript.

## Conflict of interest

PTP received speaker honoraria by Novo Nordisk. TDM received funding from Novo Nordisk. PK received speaker honoraria from Novo Nordisk, Rhythm pharmaceuticals, Hexal, Sandoz. All other authors declare that they have no conflict of interest related to this study.

## Data Deposition

Raw fastq files of the snRNAseq and further material, including statistical analyses and data, are available from the corresponding author upon reasonable request.

## Acknowledgements

We thank Miriam Krekel for technical assistance and assistance with animal studies. RGA wants to thank the Universidad Nacional Autónoma de México and Dirección General de Asuntos del Personal Académico (UNAM/DGAPA/PASPA) for the stipend given during the sabbatical year. Figures 1 and 2 were created with BioRender.com.

## Funding

This work was funded by the Deutsche Forschungsgemeinschaft (DFG, German Research Foundation) - Project-ID 424957847 - TRR 296 (AM, PK, BO, RO, TDM, PTP), and in part by the European Research Council ERC-CoG Yoyo-LepReSens (no. 101002247), awarded to PTP (RGA, CM, NM, MB). MB received funding from the Helmholtz International Research School for Diabetes graduate program. TDM received funding from the DFG (TRR296, TRR152, SFB1123 and GRK 2816/1) and the European Research Council ERC-CoG Trusted no. 101044445. PK received funding from the DFG (KU 2673/6-1, KU 2673/7-1) and the European Research Council ERC-CoG E-VarEndo 101043991.

## Supplemental Figures with figure legends

**Supplemental Figure 1:**
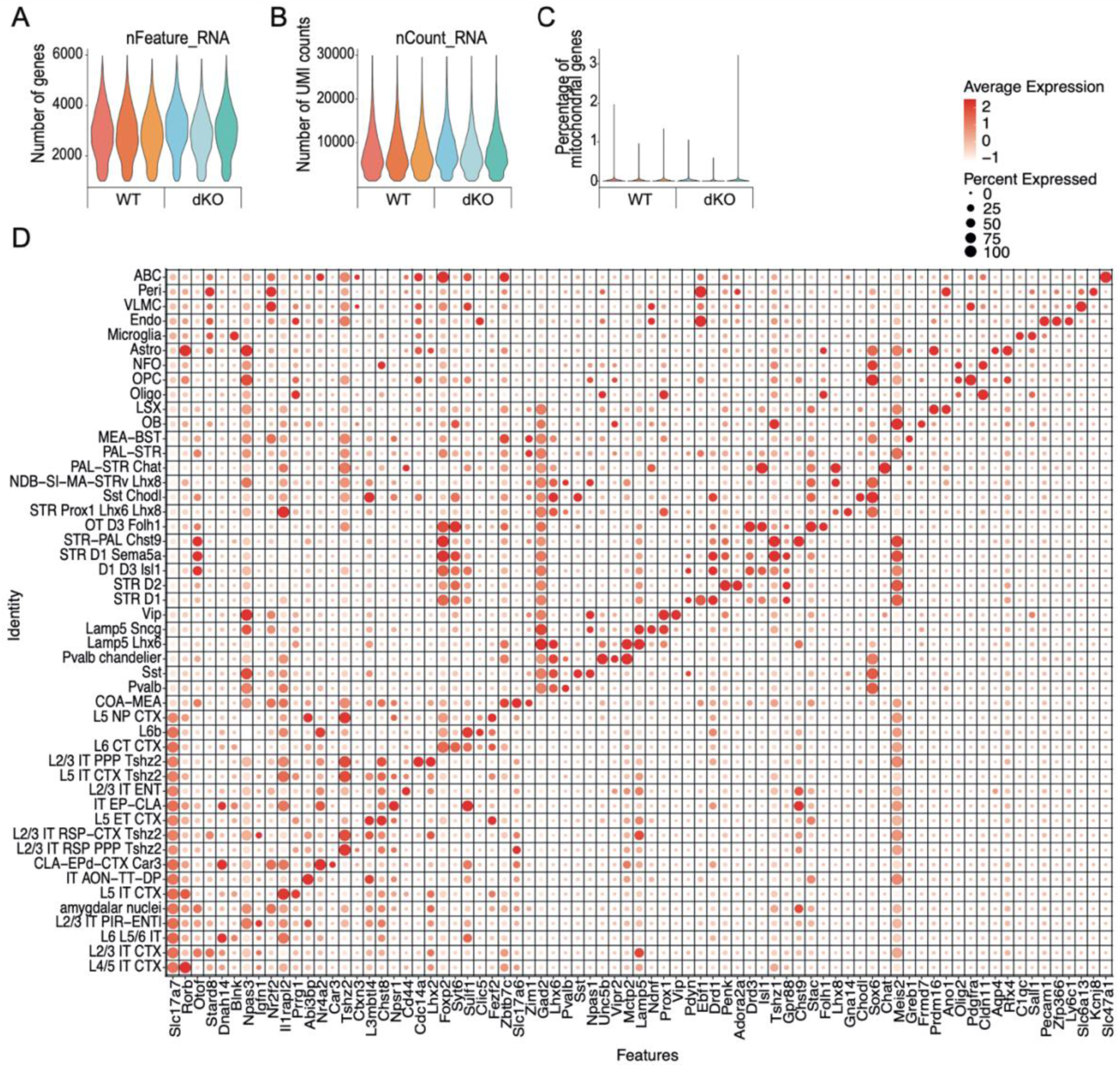
Quality assessment of nuclei from MCT8/OATP1C1 dKO mouse cortices passing the snRNAseq pipeline and marker gene patterns for the respective clusters. The nuclei passed quality checks for (A) average genes, (B) unique molecular identifier (UMI) reads per nucleus, and (C) the lack of mitochondrial genes. (D) Dotplot showing expression of some marker genes (x-axis) for the 48 clusters (y-axis). N = 3 per genotype.

**Supplemental Figure 2:**
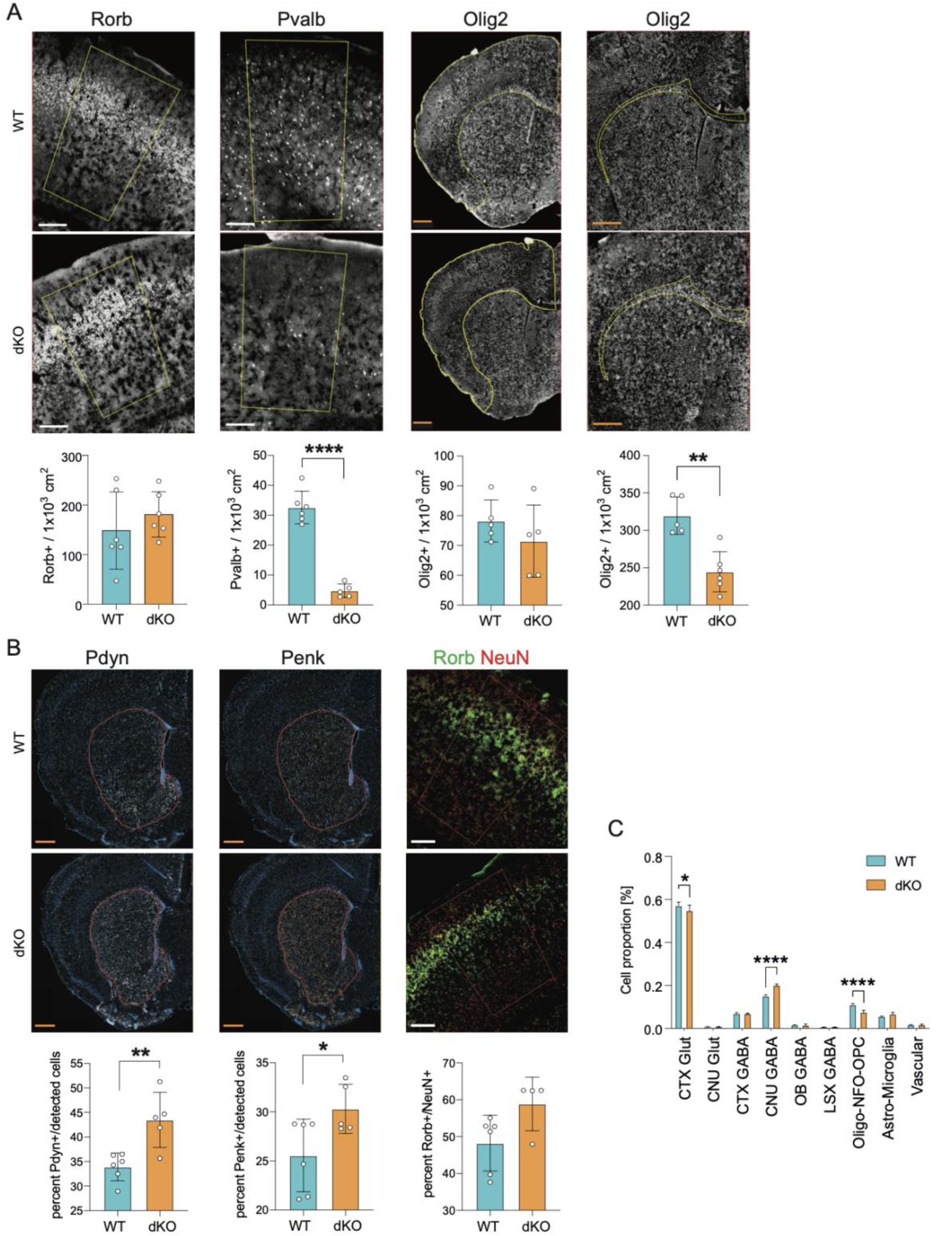
Validation of changes in proportional composition of neuronal and non-neuronal cell types. (A) Immunofluorescent stainings for Rorb, as marker for cluster L4/5 IT CTX, Pvalb, and Olig2, as marker for the Oligo cluster in WT and dKO mouse brains at P21. Cells positive for the immunostaining were counted using QuPath [1] and normalized to the area of annotation marked in yellow, for Olig2 either the entire cortical area or only the corpus callosum. (B) RNAscope analysis for Pdyn, Penk, and Rorb expressing cells. RNAscope+ cells were counted in the annotated area and normalized to the number of cells stained with DAPI (blue) for Pdyn and Penk, or to NeuN+ neurons for Rorb. (C) Cell proportion differences of the overarching groups between WT and dKO. (A-B) White scalebar = 200 µm, orange scalebar = 500 µm. Values represent means ± SD. N = 4-6 per genotype. Statistical significance was determined using one-tailed unpaired Students’ t-tests with Welch’s correction. * p < 0.05, ** p < 0.01, and **** p < 0.0001. (C) N = 3 per genotype. Statistical significance was determined using multiple t-tests with Holm-Sidak correction. * p < 0.05, **** p < 0.0001.

**Supplemental Figure 3:**
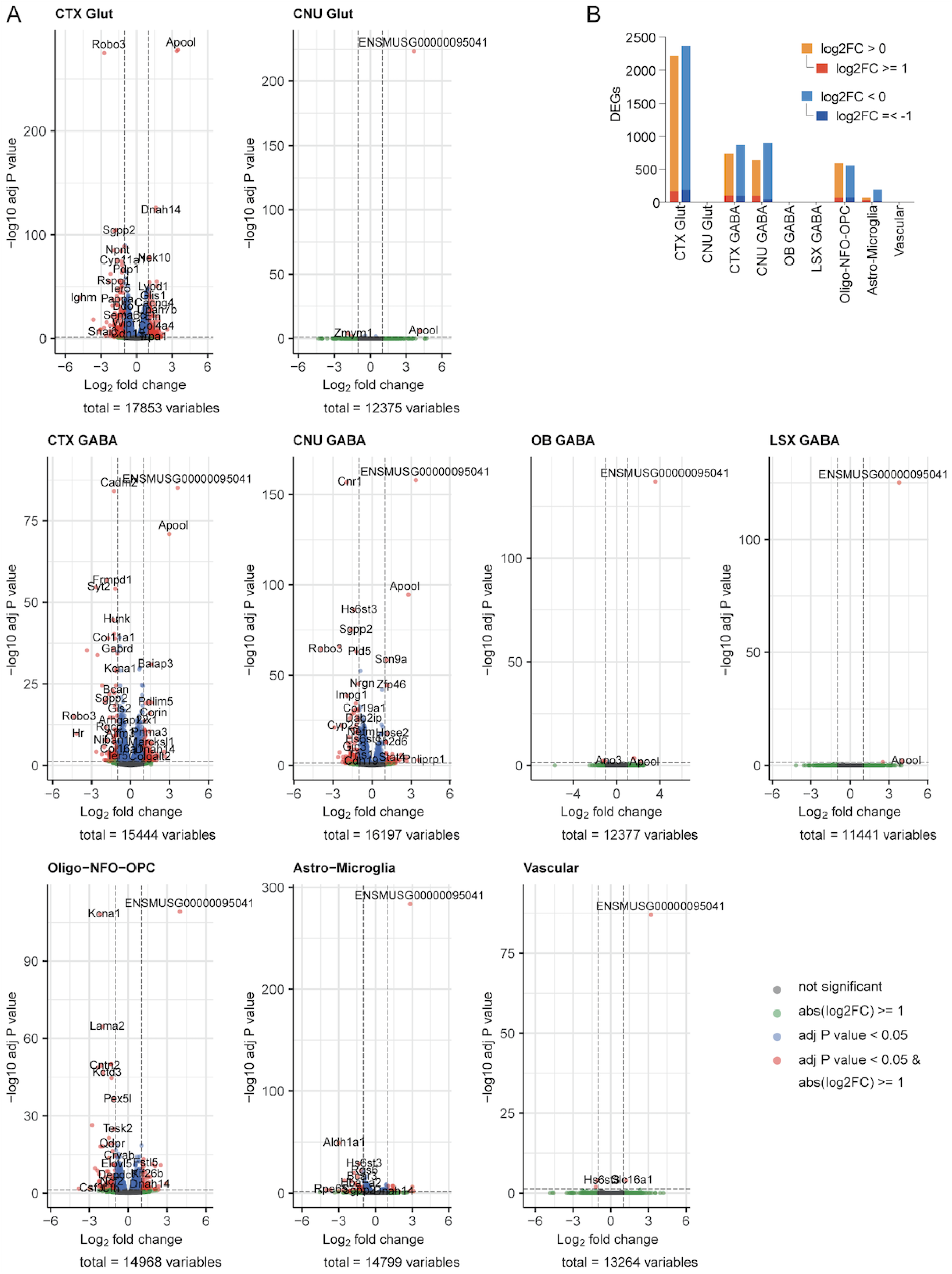
Differential gene expression in the overarching glutamatergic, GABAergic, and non-neuronal cell groups. (A) Volcano plots per overarching cell type. (B) DEGs per overarching cell type with padj < 0.05, those upregulated in dKO in red/orange and downregulated in blue/light blue. Red and blue are genes that additionally have a log2FC greater than 1 or less than −1.

**Supplemental Figure 4:**
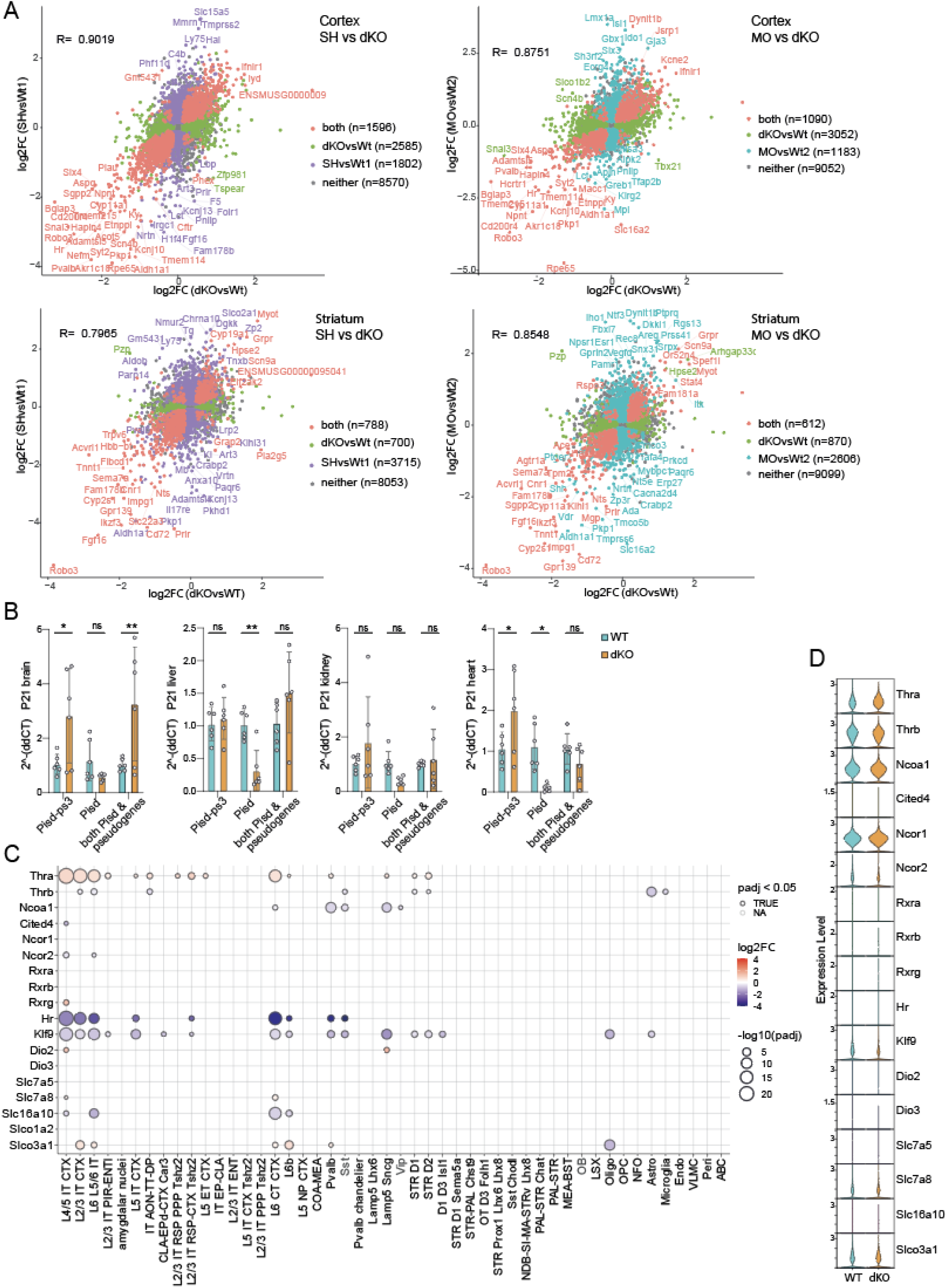
Correlation to the published SH and MO bulkseq data and expression of thyroid hormone effectors. (A) Correlation of DEGs in the cortex or striatum bulkseq datasets of SH or MO with the cortex or striatum pseudobulk version of our dKO dataset, respectively. DEGs significant in both SH and our dKO or significant in both MO and our dKO, respectively, are plotted in orange (“both”). DEGs significant only in our dKO are plotted in green, significant only in SH in purple, significant only in MO in blue, or not significant (padj >= 0.05, “neither”) in grey. Genes are plotted with their log2FC values in the SH or MO datasets on the y-axis and the log2FC values in our pseudobulk datasets on the x-axis. Correlation based on genes with padj < 0.05. (B) qPCR for *Pisd-ps 3* (ENSMUSG00000095041), actual *Pisd*, and for a common region of *Pisd* and its pseudogenes in brain, liver, kidney, and heart from 21d old WT and dKO mice. Values represent means ± SD. N = 6, multiple Students’ t-tests with the Holm-Sidak method. * p < 0.05, ** p < 0.01. (C) Known effectors of thyroid hormone action and their log2FC between WT and dKO clusters, or (D) their expression values between WT and dKO in general.

**Supplemental Figure 5:**
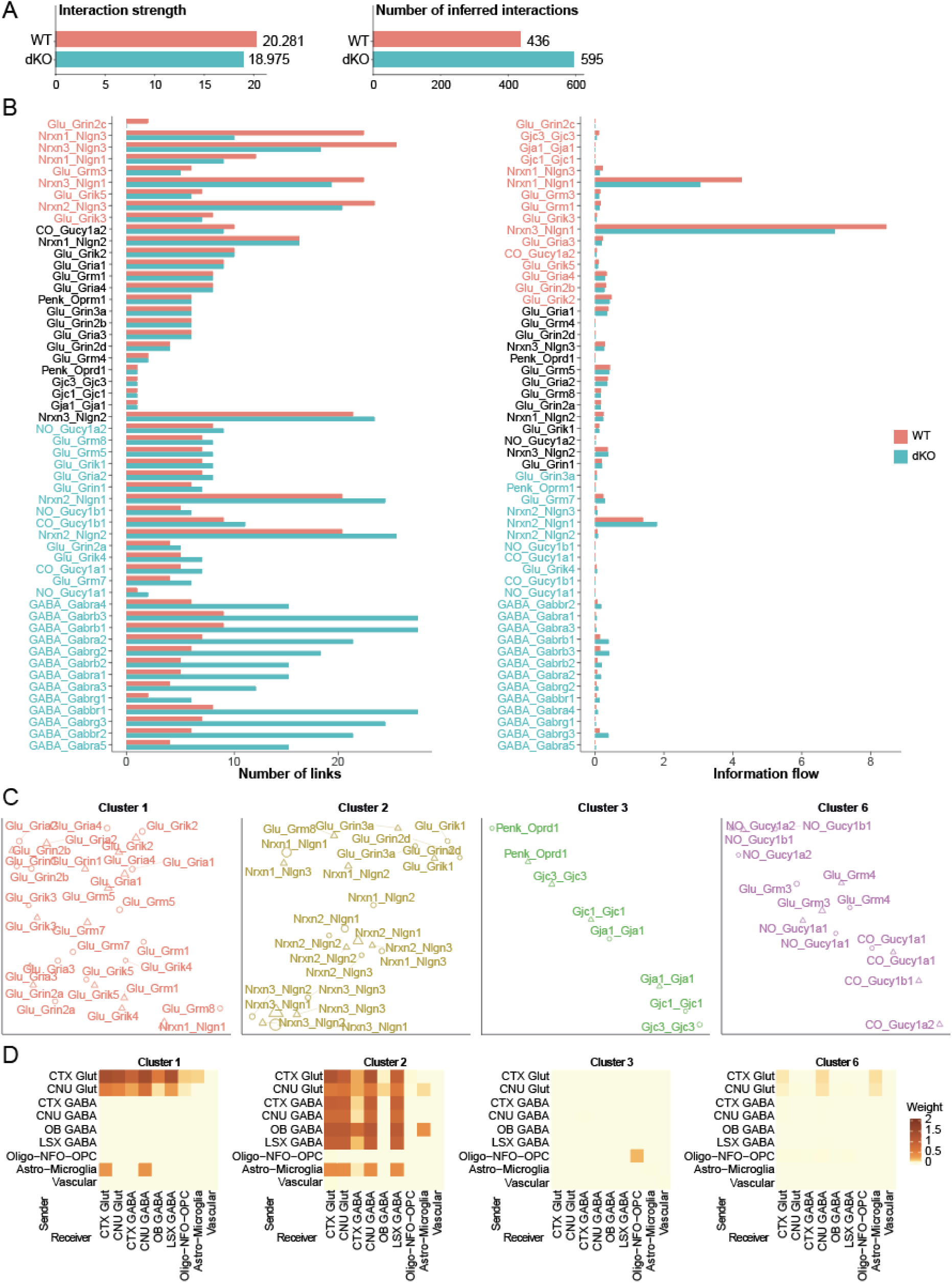
Cell-cell communication profiling in overarching cell groups of WT and dKO mouse cortices using NeuronChat. (A) Interaction strength and number of interactions inferred using NeuronChat. (B) Number of links (left) and the quantitative information flow (right) for specific interaction-pairs in WT vs. dKO mice. Bars are sorted and interaction pairs are colored by their ratio between WT and dKO. (C) UMAP of clusters 1 to 3 and 6 corresponding to Figure 6 (C) and (D) heatmaps depicting the weight/strength of communication signals for these interaction-pair clusters.

**Supplemental Figure 6:**
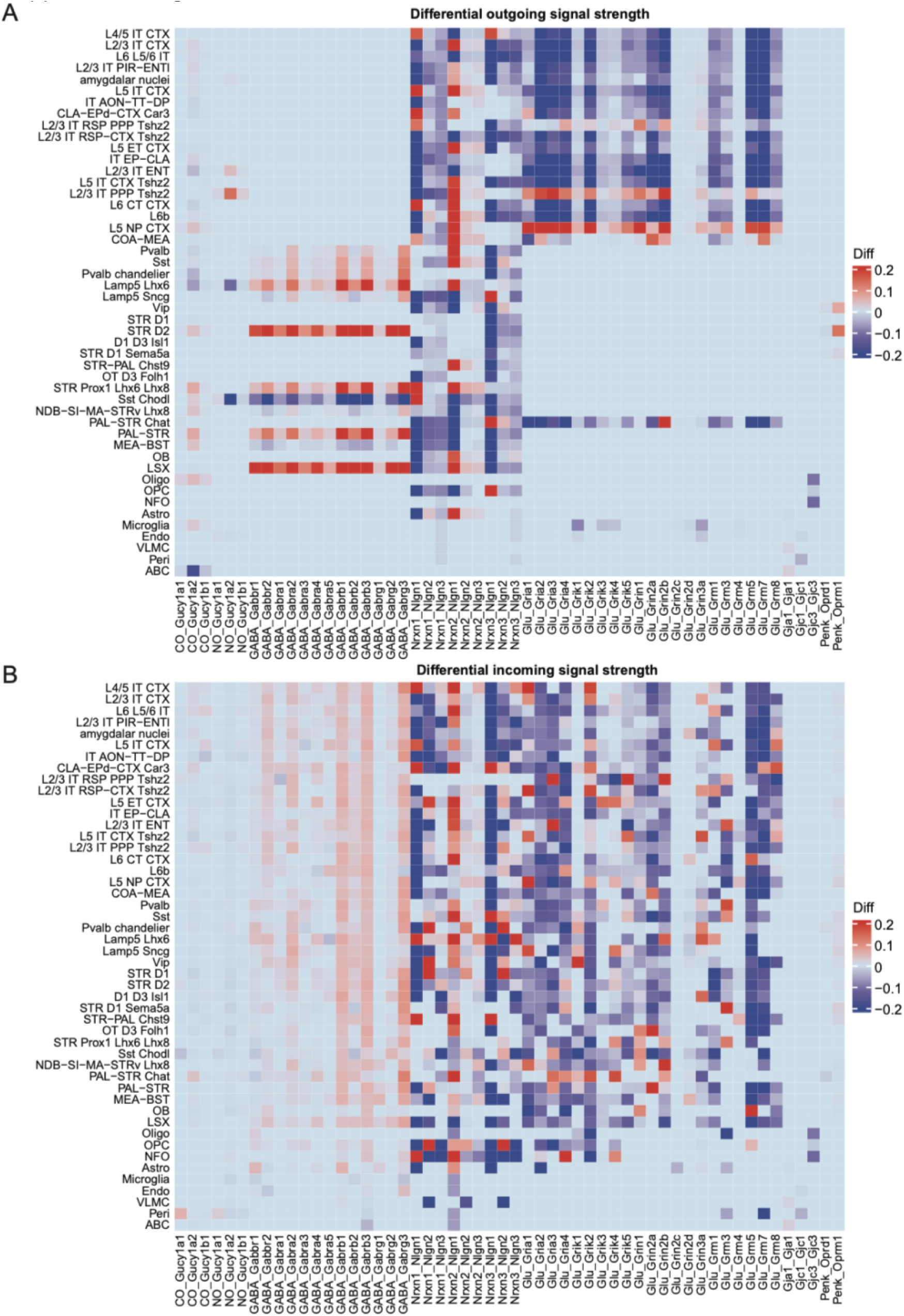
Neuron-Chat analysis of differential outgoing and differential incoming signal strengths of specific interaction pairs in individual cell clusters. Heatmaps of (A) differential outgoing and (B) differential incoming signal strength of specific interaction pairs (x-axis) for WT vs dKO mice between individual cell clusters using NeuronChat.

**Supplemental Figure 7:**
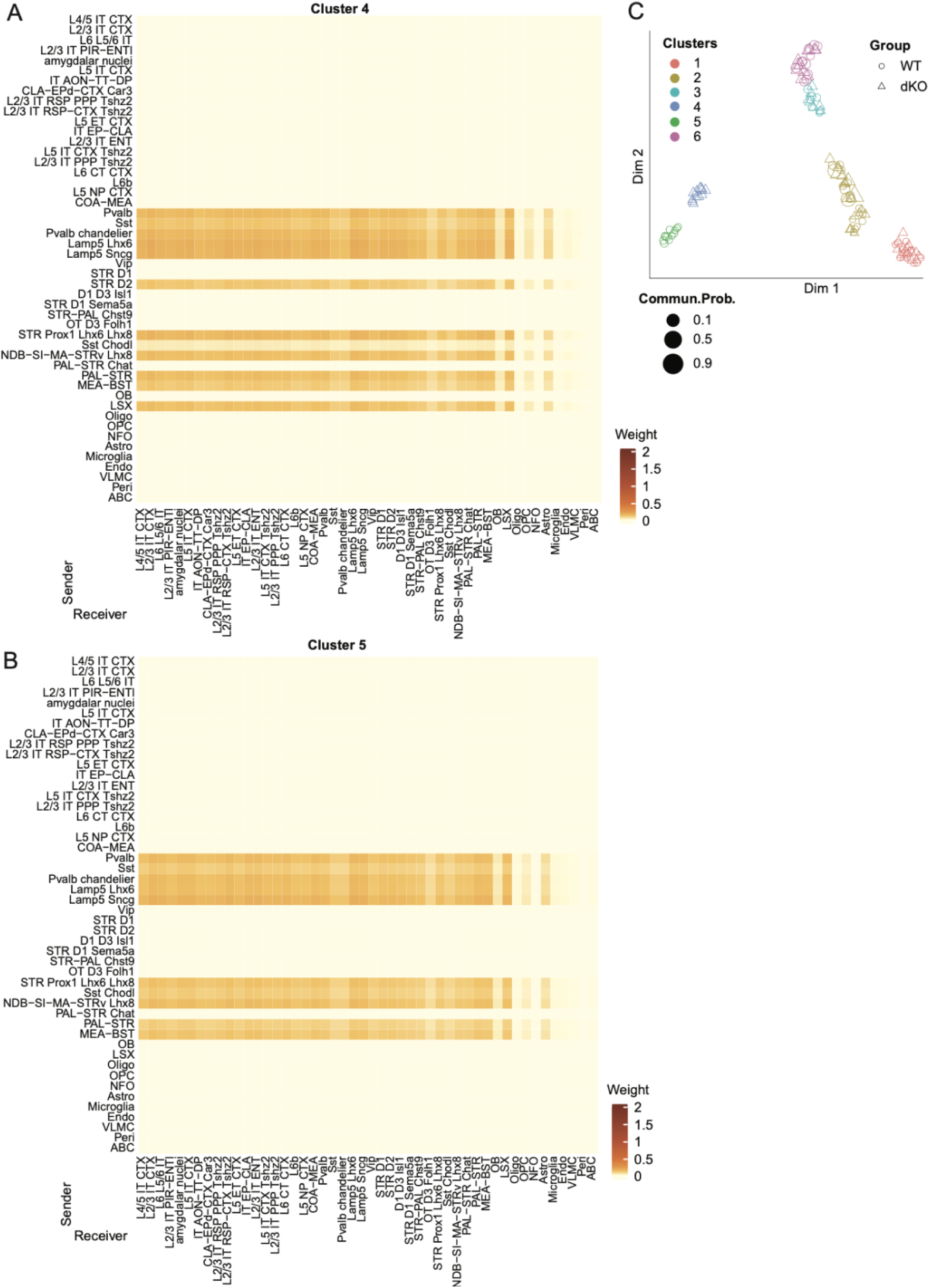
Unique, cell-type specific GABAergic communication networks in WT and dKO mice. Consistent with data obtained from to the overarching cell groups, Neuron-Chat based cell-cell-communication analyses of interaction pairs for individual cell types reveals two distinct clusters based on sender-receiver similarities that were (A) unique for dKO (Cluster 4) and (B) WT (cluster 5) mice. The corresponding heatmaps depict the weight/strength of communication signals between the individual clusters as senders and receivers for specific interaction-pairs. The respective weights of signal in the heatmaps represent the sum of communication strength values over all interaction pairs of that cluster. (C) Two-dimensional manifold projection of the 6 overall clusters and the unique communication networks for cluster 4 (dKO) and cluster 5 (WT). Statistical significance was assessed with NeuronChats integrated statistical tests, with p-values corrected for multiple tests using the Benjamini-Hochberg procedure.

**Supplemental Figure 8:**
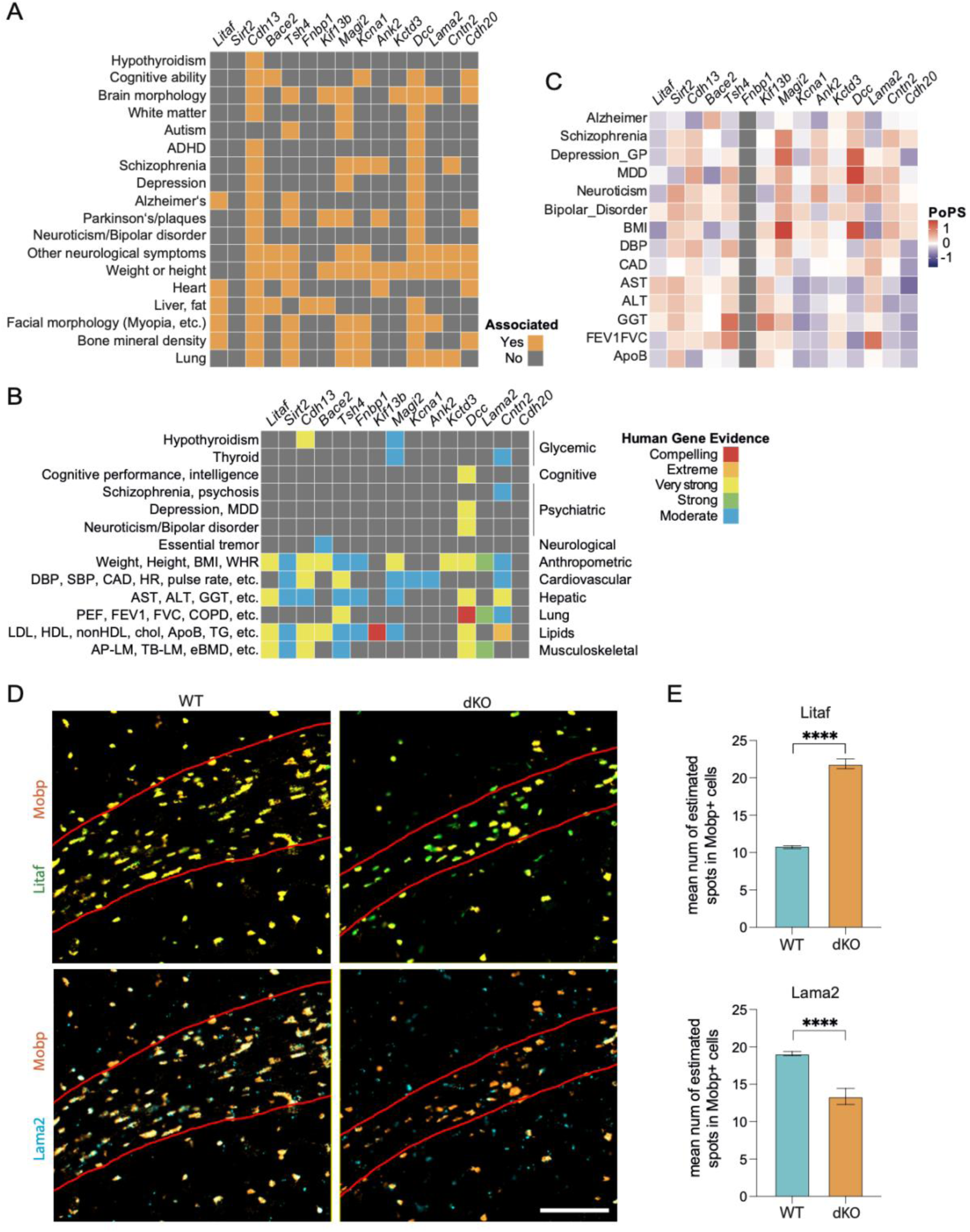
Human genome wide association of genes differentially expressed between WT and dKO oligodendrocyte subclusters to AHDS phenotypes. (A) Selected traits from the NHGRI-EBI GWAS catalogue related to AHDS symptoms. Marked orange are the oligodendrocyte related genes (Fig. 7 F-G) reported to be associated with the trait. (B) Summary of phenotypes from the human gene evidence. The evidence for the genes’ association with the phenotypes (left) was ranked from compelling to moderate. The phenotype’s group is noted on the right. (C) Polygenetic priority score, showing which genes compiled further evidence for involvement in the phenotypes. (D) Expression of two selected genes, Litaf and Lama2, in the WT and dKO P21 mouse brain at the corpus callosum using RNAscope. Mobp was used as oligodendrocyte marker. Scalebar = 100 µm. (E) Quantification of Litaf and Lama2 expression using the mean number of estimated spots in Mobp+ cells generated using QuPath. Statistical significance was determined using one-tailed unpaired Students’ t-tests with Welch’s correction. Plotted as mean with SEM. N = 4-6 per genotype of which each cell was treated as individual value. **** p < 0.0001.

